# Structural diversity of Arc oligomers within the excitatory synapse

**DOI:** 10.1101/2024.06.05.597462

**Authors:** Martina Damenti, Giovanna Coceano, Mariline Mendes Silva, Jonatan Alvelid, Chiara Sgattoni, Andrea Volpato, Lea Rems, Luciano A. Masullo, Eduard M. Unterauer, Rafal Kowalewski, Lucie Delemotte, Erdinc Sezgin, Ralf Jungmann, Ilaria Testa

## Abstract

The Activity-Regulated Cytoskeleton-Associated protein (Arc), is pivotal to mediate plasticity responses in neuronal cells. *In vitro* studies suggest its ability to form high- and low-order oligomers, which are involved in neuronal trafficking. Despite its important functions, no direct observation of Arc oligomers in cells has been presented due to its highly regulated spatiotemporal expression, the small size of the structures, the lack of appropriate labelling strategies and the background associated to free diffusing cytosolic proteins. Here, we apply super resolution microscopy to observe Arc oligomeric states in cellular environment with focus on the excitatory synapse. In cells, we provide the first evidence of Arc high-order oligomers; we uncovered intermolecular interactions of Arc, its tendency to form liquid condensates and interaction with lipid bilayers. Arc high-order oligomers affect AMPA receptor surface levels. Together, our observations suggest a model by which Arc oligomerization mediates plasma membrane negative curvature favoring AMPA receptors endocytosis.

## Introduction

Arc is an immediate early gene transcribed upon neuronal activity^1–4^. Its local translation in dendrites is crucial to mediate plastic responses and fine tune the nanosized molecular organization of synapses to ultimately ground learning and memory^5,6^.

Arc engages in the endocytosis of ionotropic AMPA receptors (GluA) to mediate homeostatic synaptic scaling and long-term depression. It was found to interact with several proteins involved in endocytosis, including Endophilin-3 and Dynamin-2^7^, as well as with GluA auxiliary subunit Stargazin (Tarpγ2)^7, 8, 9, 10^. Additionally, Arc influences the transcription of GluA1 subunit within the nucleus^11^. On the contrary, in long-term potentiation Arc stabilizes the actin-cytoskeleton of dendritic spines^12, 13^. The seemingly antagonistic role in synaptic plasticity puzzled neuroscientists until the observation of Arc’s tendency to oligomerize. Phylogenomic analyses reported the homology of tetrapod Arc gene to Ty3-Gypsy retrotransposon family. *In vitro* experiments showed Arc forming structures ranging from dimers, to 32-mers^14–18^, and even up to retroviral capsid-like particles comprising approximately 130 units^17, 19–21^. Furthermore, Arc was also found in exosome-sized extracellular vesicles, released and taken up by recipient neurons ^17, 22^.

Hence, the presence of distinct Arc *n*-meric states may be necessary to support functions, such as receptors endocytosis, cytoskeletal binding or viral-like interneuronal transfer. In different heterologous systems, we^23^ and Hedde et al.,^24^ have demonstrated that Arc forms, together with monomeric and low-order oligomers, rigid nanometric particles. In primary cortical neurons and in rat dentate gyrus^25^, Arc was reported to form low-abundance dimers and low-order oligomers (trimers, tetramers), which increase upon plasticity induction. Moreover, the Arc phosphomimetic mutant that impedes the formation of 32-mers *in vitro*, hinders long-term depression^18^. In neurons, however, the direct observation and quantification of various Arc *n*-meric states and their functional implications have not yet been attained, primarily due to the complex spatiotemporal regulation of Arc and its numerous interaction partners^26^.

Here, we provide the first evidence of Arc low and high order *n*-meric states organizations in neuronal cells, such as fast-freely diffusing low-orders oligomers, low-orders oligomers located within the post-synaptic density, and rigid semi-regular structures of about 60-80 nm in size, which mostly populate the endocytic zone.

Arc high order oligomerization promotes plasma membrane interaction resulting in lipid bilayer bending. A fine structural investigation with mutagenesis unraveled the Arc molecular sites responsible for the formation of high order oligomers. This enables us to control Arc oligomerization directly in neurons, which in turn affects AMPA receptors’ surface level. The high local concentration of Arc oligomers induces liquid-liquid phase separation, which results in the formation of condensed droplets comprised of freely diffusing Arc molecules. Taken together, our observations suggest a model in which Arc oligomers are recruited to the endocytic zone of the post-synaptic compartment, where they mediate plasma membrane curvature favoring AMPA receptors endocytosis. The detection of such structures, only possible with super resolution imaging, paves the way for future studies focusing on Arc synaptic release.

## Results

### Arc is organized in nanoclusters in primary neuronal cultures

According to previous studies, Arc molecules have been found in several neuronal compartments including the nucleus, soma, dendritic shafts and spines^7, 26, 27^. However, the spatial resolution of conventional fluorescence microscopy did not allow the detection of potential high- and low-order oligomers, resulting in Arc homogenously spread across the cell. Here, we used STED and 3D DNA-PAINT super resolution imaging^28^ to unravel the Arc nanoscale organization in primary neurons.

We used rat primary cortical neurons at a developmental stage (DIV6-9) where the minimal to null endogenous expression of Arc limits the observation mostly to the labelled protein. Different Arc labelling strategies were developed (Fig. 1A) to minimize the steric hindrance of the tag on potential oligomerization: Arc full-length (Arc-FL) with a SNAP-tag at the protein C-terminal^29^ (Arc-FL-C-SNAP), or between its NTD, a double α-helical domain, and the flexible linker which connects to its bilobed CTD^23^ (Arc-FL-IM-fusion-SNAP). Arc-FL-C-SNAP or Arc-FL-IM-fusion-SNAP were used, upon dye-conjugated SNAP-ligand staining, to visualize Arc in live neurons and assess the presence of Arc oligomers in absence of fixation artifacts, such as clusterization^30^. To image fixed samples, Arc was tagged either with AlfaTag^31^ (Arc-FL-C-AlfaTag, Fig. 1A, D) or mEGFP (mEGFP-N-Arc-FL^26^) followed by immunostaining (Fig. 1A, G). Endogenous Arc detection via anti-Arc antibody was disregarded for *n*-meric states quantification given the polyclonality of the most used and commercially available antibody.

**Fig. 1.**
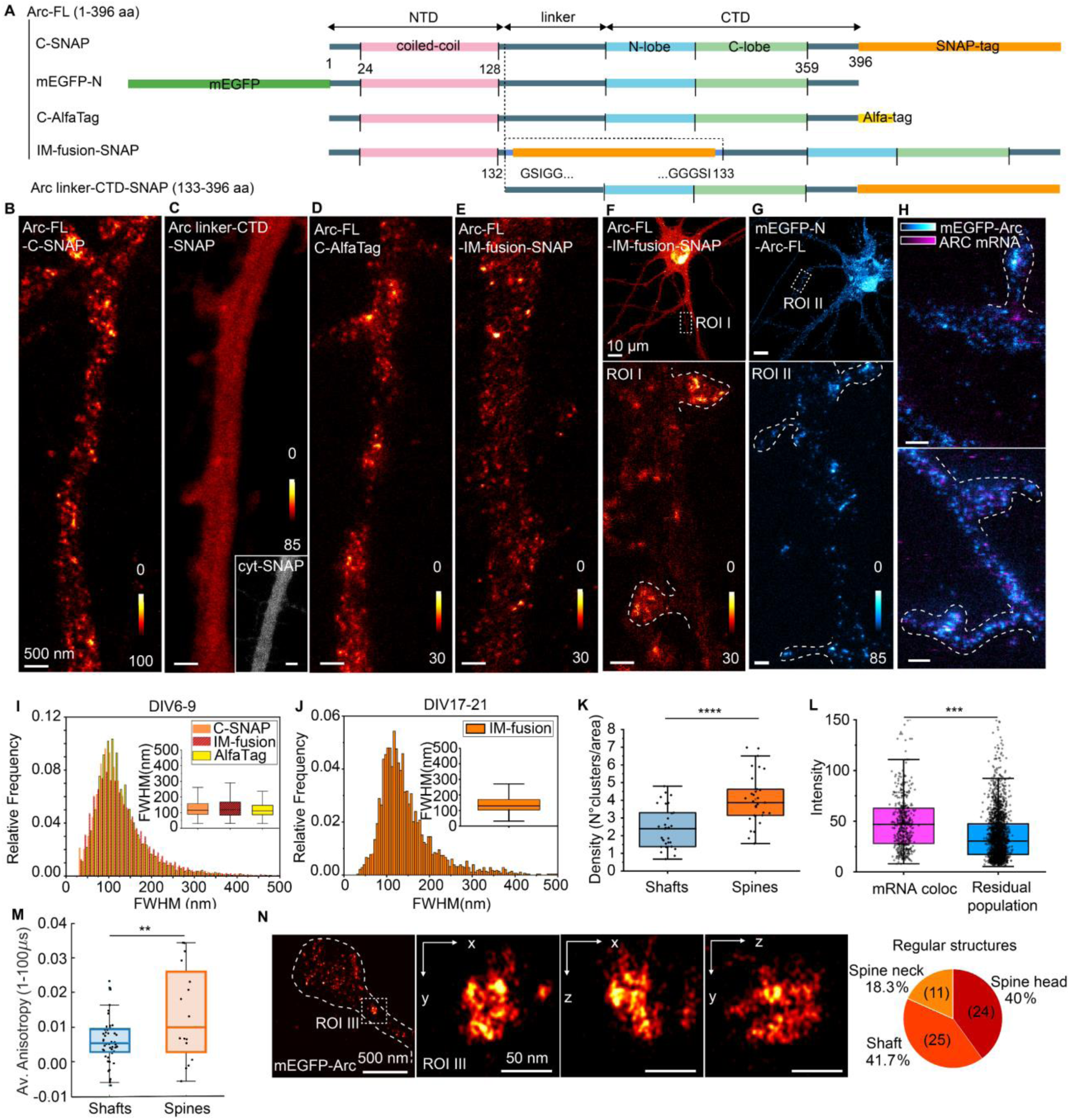
Arc is organized in nanoclusters in primary neuronal cultures. (A) Schematic representation of the Arc tagging strategies (SNAP-tag, orange; AlfaTag, yellow; mEGFP, green) and positioning (N-term, C-term, IM-fusion) used in the work for Arc-FL (1-396 aa) and linker-CTD truncated mutant (133-396 aa). Arc domains are in different colors (NTD, pink; flexible linker, black; N-lobe, blue; C-lobe, green). (B) Example of a cortical neurite (DIV6-9) expressing Arc-FL-C-SNAP in live STED microscopy. (C) Example of a cortical neurite (DIV6-9) expressing Arc linker-CTD-SNAP, where no nanoclusters could be identified in live-cell STED microscopy, similarly to cytosolic SNAP (cyt-SNAP, inset). (D) Example of a neurite (DIV7) expressing Arc-FL-C-AlfaTag in STED microscopy. (E) Example of a cortical neurite (DIV6-9) expressing Arc-FL-IM-fusion-SNAP where the tag is placed as intramolecular fusion (IM-fusion) between the NTD and the flexible linker in homology with human HIV-1 GAG protein. (F) Mature Primary Neuron (DIV21) genetically encoding Arc-FL-IM-fusion-SNAP. In ROI I Arc nanoclusters show localization in dendritic shaft and in the spines’ head. (G) Mature Primary Neuron (DIV21) genetically encoding mEGFP-N-Arc-FL. In ROI II, Arc nanoclusters show localization in dendritic shaft and in the spines’ head. (H) Mature Primary Cortical Neuron (DIV20) genetically encoding mEGFP-N-Arc-FL (cyan) where Arc mRNA (magenta) was labelled via RNA FISH. A subset of Arc nanoclusters is shown to colocalize with the mRNA. (I) The histogram reports Arc nanocluster FWHM distributions from three labelling strategies. Arc-FL-C-SNAP (22 DIV6-9 cortical neurons from 5 independent cultures, N=24357 fitted nanoclusters). Arc-FL-IM-fusion-SNAP (13 DIV6-9 cortical neurons from 3 independent cultures, N=12429 fitted nanoclusters). Arc-FL-C-AlfaTag (10 DIV7 hippocampal neurons, N=4282 fitted nanoclusters). As an inset, the box plot reports the difference in the Arc nanoclusters FWHM distribution for three labelling strategies. (J) The histogram reports Arc-FL-IM-fusion-SNAP nanocluster FWHM distribution (18 (DIV17-21cortical neurons from 5 samples in 2 independent cultures, N=24357 fitted nanoclusters). (K) The box plot represents the significantly higher density of Arc nanoclusters in spines (orange) versus dendritic shaft (blue). Each data point is the average density value per spines and shaft per image (18 DIV17-21 cortical neurons from 5 samples in 2 independent cultures, N=24357 fitted nanoclusters, two-sample two-sided Student’s t test p-value= 1.11×10^-4^, box plots show the 25–75% interquartile range, with the middle line representing the mean, and the whiskers derived from 1.5 * interquartile range). (L) Box plots showing the significantly higher intensity of Arc nanoclusters in colocalization with Arc mRNA (two-sample two-sided Student’s t test p-value=1.81×10^-15^). Data are obtained from 4 cortical neurons (DIV20) (N=1075 fitted nanocluster). Box plots show the 25–75% interquartile range, with the middle line representing the mean, and the whiskers derived from 1.5 * interquartile range). (M) Average fluorescence anisotropy in the time window 1-100 µs measured with STARSS for Arc-IM-fusion-rsEGFP2 in spines (orange) and in shafts (blue) from 3 independent cultures in four sessions in different days. Shaft anisotropy from 50 FOVs: 0.0061± 0.0020 CI 95%, median: 0.005, quartiles: 0.003 – 0.009, whiskers: -0.006 – 0.016; spines anisotropy from 16 FOVs: 0.0139 ± 0.0072 CI 95%, median: 0.0099, quartiles: 0.003 – 0.026, whiskers: -0.0056 – 0.034; T-test p-value = 0.003. (N)(left panels) Example of a dendritic spine from cortical neurons (DIV22) expressing mEGFP-N-Arc-FL imaged with 3D DNA-PAINT. ROI III shows an example of Arc nanocluster with radially symmetric structures shown in x-y, x-z and y-z orientation. (right panel) Pie chart showing the localization of 60 Arc nanoclusters (21 neurons from 2 independent cultures). localized in the head of dendritic spines (40%, N=24), in the spine neck (18.3%, N=11) and in dendritic shafts (41.7%, N=25).

Regardless of the labelling strategy, STED microscopy showed Arc localized in the nucleus (Fig. S1A), in the cytosol (in live samples) and in a broad range of nanoclusters sizes (30 to 300 nm FWHM), peaking at 95-105 nm (Fig. 1I). The observed nanoclusters were not induced by artificial tag clusterization but by Arc oligomerization. Indeed, the expression of SNAP-tagged truncated mutant linker-CTD (Fig. S1H), which lacks the NTD required for Arc oligomerization^17,14^ showed a homogenous cytosolic distribution and no nanoclusters, akin to the pattern observed with SNAP-tag volume staining (Fig. 1C and inset).

To study the role of Arc nanoclusters in synaptic activity, we expressed Arc-FL-IM-fusion-SNAP in mature primary cortical and hippocampal neurons (DIV17-23, Fig. 1F). Arc nanoclusters, in a distribution peaking at 117 nm (Fig. 1J), were observed in dendritic shafts, and as denser in spines (Fig. 1K). A comparable clusters size distribution and density was confirmed in fixed samples (Fig. 1G, Fig. S1B-C).

A small population (23.2% of the total) of Arc nanoclusters was found in colocalization with Arc mRNA, detected with RNA FISH. These nanoclusters were significantly brighter than the residual population (Fig. 1H, L), and they were found also in the dendritic spines’ head.

To further investigate whether the formation of nanoclusters was affected by the high labeling densities, we used Selective Time-resolved Anisotropy with Reversibly Switchable States (STARSS)^23^. STARSS probes the rotational diffusivity of molecules to extract the hydrodynamic radius, and, hence, it enabled the quantification of nanoclusters sizes, including both labeled and unlabeled Arc molecules. In neurons expressing Arc-IM-fusion-rsEGFP2 (DIV16-17), the average anisotropy in the time window 1-100 µs was higher in spines compared to dendrites (Fig. 1M), suggesting that Arc tended to form larger nanostructures when located in spines. The observed anisotropy decay, averaging from 0.014 - 0.006 within the observed temporal window, was compatible with the presence, along with a fast diffusive component of monomers or low-order oligomers, of a fraction of Arc molecules, at least 15% of the total fluorescently labelled protein, anchored to structures of roughly 60-90 nm in diameter. Simultaneously, a smaller fraction of the protein, roughly half of the 60-90 nanometer fraction, was involved in larger structures, greater than 200 nm or static (Fig. S1D).

Given the broad size distribution of Arc nanoclusters observed in STED microscopy, we applied DNA-PAINT to further inspect their potential suborganization. Primary neurons expressing mEGFP-N-Arc-FL imaged with 3D DNA-PAINT, revealed the presence of different organization: Arc low-order oligomers located in the cytosol, which presumably were the fast monomeric and low-order components measured in STARSS, Arc low-order oligomers accumulated within shafts and spines domains, and an additional type of high-order oligomers of about 60-80 nm in diameter (26.6 x 10^4^ nm^3^ on average) with a tendency to radial symmetry (Fig. 1N). These structures were preferentially localized in the spines (respectively 40% in spine heads and 18.3% in spine necks), but found, as well, in the shaft (41.7 %) (Fig. 1N, right panel). By dividing their average volume by that of the low-order oligomers (Fig. S1E, F), we estimated that the high-order structures are composed of approximately 60 copies of the low-order counterpart. Assuming that these low-order oligomers were primarily monomers, we calculated an average copy number of 60 units per structure. The overall number was relatively rare (N = 64 in 21 neurons from 2 independent experiments), but it did not correlate with mEGFP-N-Arc-FL expression level of the cell, indicating that they were not artifacts of the protein overexpression (Fig. S1G).

We further characterized which part of molecules is responsible for Arc oligomers formation with a specific focus on higher *n*-meric states (Fig. S1I-M). Our spectroscopic and imaging data on different Arc truncation mutants expressed in HeLa cells suggested that both NTD-NTD and NTD-CTD intermolecular interactions are involved in *n*-merization. While the NTD-NTD interaction drives the oligomerization up to a tetramer, we propose that CTD-NTD interaction, which prevents NTD-NTD tetramerization, is relevant for oligomerization steps above the tetrameric state. The data are well supported by the Alphafold Multimer interaction predictions (Fig. S1K-M). Altogether our data demonstrated the tendency of Arc to assemble in distinct complexes which localize preferentially in dendritic spines and can contain Arc RNA.

### Arc nanoscale organization at the excitatory synapse

The observation of distinct Arc oligomers in dendritic spine heads of neurons suggests their involvement in distinct synaptic functions. In mature neurons (DIV21) endogenous and immunostained Arc appeared distributed in nanoclusters all over the cytosol, with highly variable expression levels from neuron to neuron (Fig. 2A). Considering Arc’s role in AMPA receptors’ endocytosis, its interaction with PSD95 and its function as an actin stabilizer during long-term potentiation, we characterized Arc’s organization at both the synaptic and endocytic zone (EZ) level. We defined the post-synaptic endocytic zone as an area located within 200 nm from PSD95 perimeter ^39^. Arc nanoclusters at the synaptic and EZ were significantly larger and brighter than the extra-synaptic population (Fig. 2D, E and Fig. S2A; synaptic Arc median FWHM: 92 nm, EZ Arc median FWHM: 79 nm, extra-synaptic Arc median FWHM: 73.5 nm). Another way to look at the endogenous Arc population was to express mEGFP-N-Arc-FL under the control of Arc endogenous promoter, as previously reported by Okuno et al., 2012 ^26^. STED imaging showed Arc in proximity and in colocalization with the immunostained PSD95 (Fig. 2B). Synaptic Arc nanoclusters displayed larger diameter and increased brightness than the extra-synaptic ones (Fig. S2B, C), likewise to the immunostained counterpart.

**Fig. 2.**
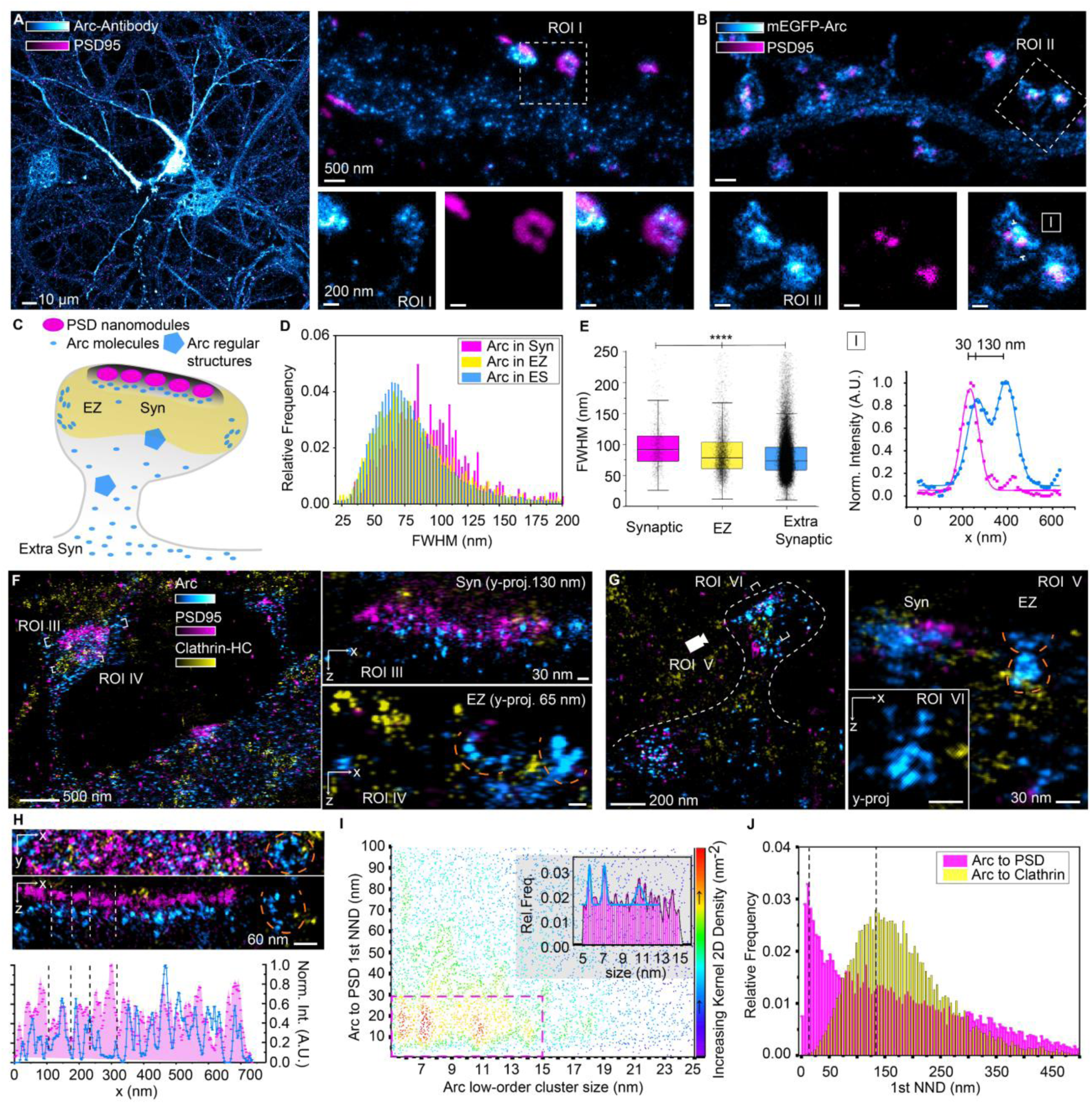
Arc nanoscale organization at the excitatory synapse. (A) Representative image of primary cortical neurons (DIV21) with Arc (cyan) and PSD95 (magenta) immunostained. ROI I shows close-up of synapses where Arc colocalizes with PSD95. (B) Mature cortical neurite (DIV21) expressing mEGFP-N-Arc-FL (cyan) under the control of Arc endogenous promoter and immunostained for PSD95 (magenta). In ROI II, Arc shows nanoclusters of different sizes localized in the head of the spine in proximity and co-localizing with PSD95. (lower panel) Line profile traced along the white arrows. (C) Cartoon representation showing the different Arc subpopulations which localize within the peri-synaptic level as observed in STED and DNA-PAINT imaging. (D) The histogram reports the distributions of Arc nanocluster sizes for Arc in co-localization with PSD95 (Syn, magenta), within the endocytic zone (EZ, within 200 nm from PSD95 perimeter, yellow) and extrasynaptic (ES, blue) (26 neurons from 3 independent cultures). (E) The box plot shows the significant difference in nanocluster sizes between Synaptic Arc in co-localization with PSD95 (Syn), Arc within the EZ (EZ, within 200 nm from PSD95 perimeter) and extrasynaptic Arc (ES). Box plots show the 25–75% interquartile range, with the middle line representing the mean, and the whiskers derived from 1.5 * interquartile range. Two-sample two-sided Kolmogorov–Smirnov test p-values respectively: 2.6934e-4, 1.6397e-17, 1.4041e-18). Data derived from 28 neurons from three independent experiments in three independent cultures. (F) Example of a dendrite with synapses located in spine heads and in the shaft from DIV22 cortical neuron expressing mEGFP-N-Arc-FL (cyan), immunostained for PSD95 (magenta) and Clathrin-HC (yellow) and imaged with 3D DNA-PAINT. Arc accumulates at the synaptic level (Syn) for certain synapses (ROI III) but not for others. At the EZ level, Arc semi-circular structures can be observed (ROI IV, dashed orange lines). (G) Example of a dendritic spine from mEGFP-N-Arc-FL expressing DIV22 cortical neuron (cyan), immunostained for PSD95 (magenta) and Clathrin-HC (yellow) and imaged with 3D DNA-PAINT. In ROI V, observed from the white camera icon direction, Arc is shown in accumulation at the synaptic level (Syn) and in semi-circular and regular structures at the EZ level (EZ). ROI VI shows a 65 nm y-projection of Arc regular structures perpendicularly sliced along white squared lines. (H) Example of a dendritic shaft synapse from DIV22 cortical neuron expressing mEGFP-N-Arc-FL (cyan) and immunostained for PSD95 (magenta) and Clathrin-HC (yellow) imaged with 3D DNA-PAINT. Observing the synapse axially (130 nm y-projection) Arc molecules are shown to partially align with PSD95 nanomodules as reported in the line. Next to PSD95 an example of Arc circular structure is observed. (I) The scatter plot shows the correlation between Arc PSD95 1^st^ NNDs to Arc low-order cluster size, the density color map reveals three higher density areas corresponding to subpopulations of Arc diameters of 5.7, 7.2 and 10.7 nm within 30 nm from PSD95. The frequency distribution in the inset reports the peaks corresponding to the three populations. (J) The histogram reports the distributions of 1^st^ NNDs from Arc to PSD95 (magenta), which peaks at 12.5 nm, and from Arc to Clathrin-coated vesicles (yellow), which peaks at 132.5 nm. (13 neurons from 2 independent neuronal cultures).

To further investigate the potential suborganization of Arc nanoclusters at the synaptic level and characterize their *n*-meric states, we performed 3D DNA-PAINT. The expression of mEGFP-N-Arc-FL was followed by immunostaining of PSD95 and Clathrin heavy chain (Clathrin-HC), which was used as EZ marker (Fig. 2F). A comprehensive analysis of the data allowed the identification of distinct Arc populations at both the synaptic and EZ level: Arc molecules as low-order oligomers were denser in the surroundings of PSD95, and Arc’s regular and semi-circular structures were located laterally to the PSD, extending into the EZ (Fig. 2F, G). Focusing on Arc low-order oligomers at the PSD level, we found that the majority (52 out of 81) of analyzed synapses contained at least 30% more Arc molecules in colocalization with PSD95 than in extra synaptic areas. Furthermore, by measuring the distance from each Arc low-order oligomers to the 1^st^ PSD95 Nearest Neighbor (1^st^ NND), we showed that Arc can be as close as 2.5 nm from PSD95 within a distribution that peaks at 12.5 nm (Fig. 2J, magenta). The axial localizations of Arc molecules revealed distinct sub-distributions: in 55% of analyzed synapses Arc molecules aligned to PSD nanoclusters (defined as PSD95 densities in 65 nm y-projections) (Fig. 2H). Correlating each Arc low-order oligomers size to its 1^st^ PSD95 NND, we identified three distinct subpopulations of Arc sizes (5.7, 7.2 and 10.6 nm) located within 30 nm from PSD95 (Fig. 2I). The smaller populations might represent Arc monomers, while the third population, which was double in size, might reflect Arc dimer. The intermediate size could result from the strategy employed in DNA-PAINT of using two anti-EGFP nanobodies to increase the labelling density. In an alternative interpretation of the results, the intermediate size of Arc might reflect Arc dimers formed through alpha-helices coiled-coil interactions, while the third population could represent Arc tetramers.

Examining the populations lateral to the PSD within the EZ, Arc was organized in semi-circular structures adjacent, yet relatively distant from Clathrin-coated vesicles. By measuring the 1^st^ NND from Arc low-order oligomers to Clathrin-coated vesicle, we showed that Arc molecules can be as close as 7.5 nm from Clathrin-HC, with a distribution peak at 132.5 nm (Fig. 2J). However, differently from PSD95, the correlation of Arc low-order oligomers size to Clathrin-coated vesicle 1^st^ NND revealed multiple subpopulations of Arc located between 100 to 200 nm. This suggests a higher complexity of Arc-Arc interactions further away from Clathrin-coated vesicles (Fig. S2D). The broadness of the distribution might result from considering Arc molecules located both at PSD and EZ levels. In the attempt to limit the observation to Arc molecules at the EZ level, we calculated the 1^st^ NND from Clathrin-coated vesicles to Arc (Fig. S2E). In this case, the distance distribution peaked at 75 nm.

Overall, we demonstrated distinct Arc populations among Arc nanoclusters, including low-order oligomers within the post-synaptic density, and higher-order oligomers and semi-circular organization at the EZ.

### Arc oligomers form liquid condensates in cells

The compartmentalization in the post-synapse of several Arc’s interaction partners such as PSD95, CamKII, and Stargazin is phase separation-based^32, 33^. Additionally, Arc is homologous to retroviruses, whose GAG proteins are sorted by lipid-based phase partition^34^; and a family of viruses, the Birnaviridae, have been shown to form liquid phase-separated viral factories^35^. Considering that liquid-liquid phase separation and oligomerization are reported to be reciprocally interconnected: oligomers facilitate liquid-liquid phase separation^36^ and liquid-liquid phase-separation favors oligomerization^37^, we hypothesized that Arc clusters might be liquid condensates.

When Arc-FL was expressed in HeLa cell, bright and cytosolic microclusters were observed (average area: 0.705 μm^2^, roundness: 0.8, Fig. S3A, C). We verified that Arc microclusters are not lipid-bound cellular compartments or stress granules resulting from protein overexpression, given that no colocalization was observed with lipids nor with the granular marker G3BP1 (Fig. S3B). Then, we incubated HeLa cells expressing Arc-FL (C-EGFP tagged) with different concentration of 1,6-hexanediol (1,6-HD), a chemical known to affect weak protein-protein and protein-nucleic acids interaction and dissolve liquid condensates^38,37^ (Fig. 3A). We observed a 1,6-HD concentration-dependent phenotypic shift: Arc clusters areas significantly decreased (Fig. 3B), with a tendency for an increase in density up to 5% 1,6-HD, whereas at 10% 1,6-HD just a few big condensates remained (Fig. 3C).

**Fig. 3.**
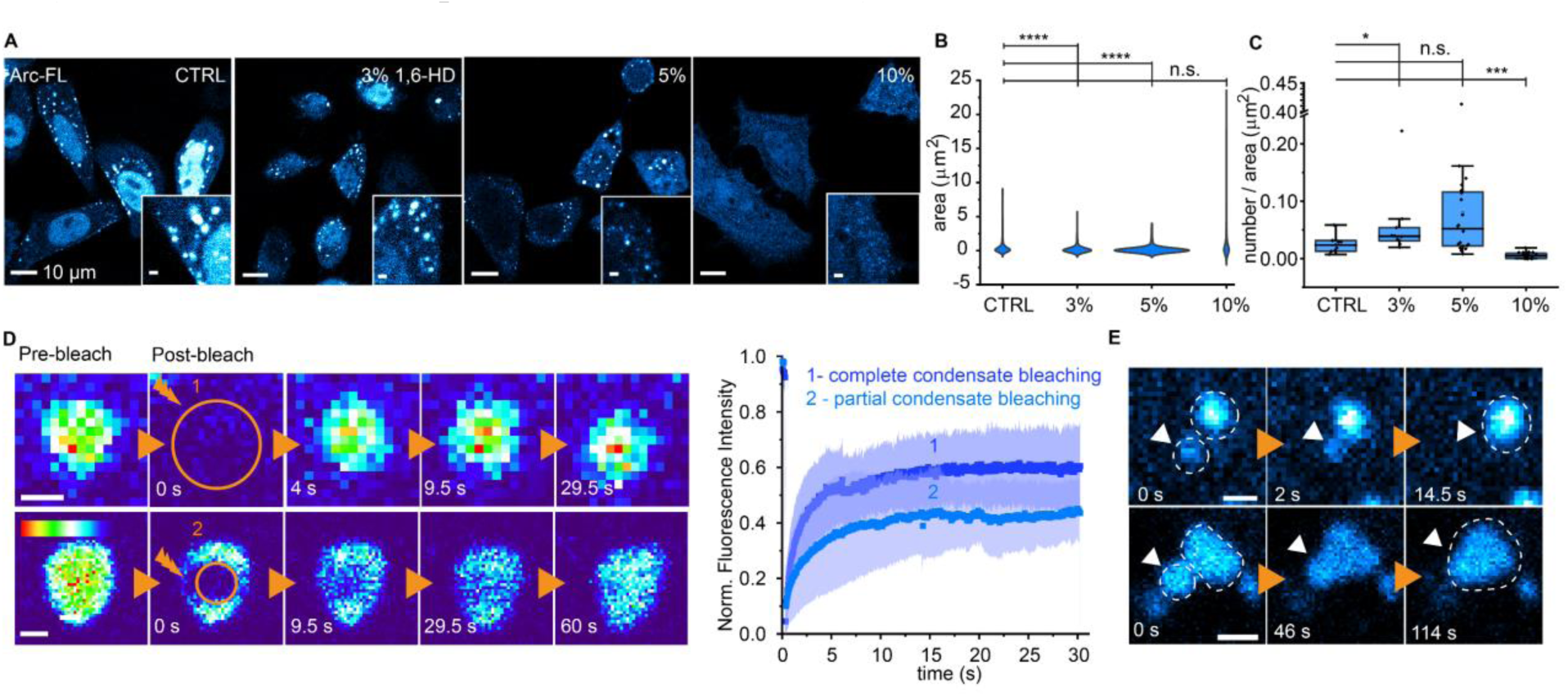
Arc oligomers form liquid condensates in cells. (A) Examples of HeLa cells exogenously expressing Arc-FL-C-EGFP untreated (CTRL) or treated respectively with 3%, 5% and 10% 1,6-HD, showing the effect of the drug on Arc condensates. (B) Violin plot showing distribution of Arc clusters area in control condition (N=10 cells) or upon treatment with respectively 3% (N=10), 5% (N=20) and 10% 1,6-HD (N=20) (Two-sample two-sided Kolmogorov–Smirnov test p-values, CTRL - 3%: p-value 1.7681×10^-28^; CTRL - 5%: p-value 1.2863×10^-48^; CTRL - 10%: p-value 0.1882. (C) Box plot showing the number of Arc condensate per area in control condition or upon treatment with respectively 3% (N=10), 5% (N=20) and 10% 1,6-HD (N=20). Box plots show the 25–75% interquartile range, with the middle line representing the mean, and the whiskers derived from 1.5 * interquartile range. Two-sample two-sided Kolmogorov– Smirnov test p-values, CTRL - 3%: p-value 0.031; CTRL - 5%: p-value 0.0946; CTRL - 10%: p-value 3.7479×10^-05^). (D)(left) Two representative time-lapses of HeLa cell exogenously expressing Arc-FL with Arc condensate fully (upper panel, 1) or partially (lower panel, 2) photo-bleached. (right) Quantitative results of FRAP for fully bleached (1, 14 clusters from 8 cells, τ_1/2:_ 2.54 (± 0.06) sec) and partially bleached (2, 35 clusters from 18 cells, (τ_1/2:_ 3.54 (± 0.05)) condensates. (E) Two representative time-lapses of HeLa cell exogenously expressing Arc-FL: condensates dynamically fuse at different time frames.

Proteins within liquid condensates are known to diffuse slower than their cytosolic counterpart, while constant diffusion in and out is maintained. To assess that, we performed Fluorescence Recovery After Photobleaching (FRAP) on Arc-FL molecules within the clusters. We completely or partially bleached the clusters to study the inter- and intra-clusters dynamics (Fig. 3D, left panel). A faster average recovery half time was measured for the completely bleached condensates, which provides evidence of a dynamic interchange of Arc molecules in and out the clusters. The immobile fraction, instead, was larger for partially bleached ones (IM_f partial_ = 68.7%, IM_f bleached_ = 51.3%) (Fig. 3D, right panel). We further assessed the mobility of the whole clusters within the sub-cellular environment with time-lapse imaging observing their tendency to dynamically fuse together (Fig. 3E). In conclusion, we have demonstrated that Arc clusters exhibit liquid-like behavior in cells. We hypothesize that these condensates could work as an additional mechanism to regulate the local availability of Arc as oligomers, keeping cytosolic concentration constant while still freely diffusing in and out at need.

### Palmitoylation and PIP lipids modulate but they are not required to mediate Arc-membrane interaction

The detection of Arc semi-circular structures at the EZ suggests potential endocytosis events. In addition, the expression of Arc-FL in HeLa cells resulted in prominent plasma membrane localization (Fig. S1I). Most notably, the plasma membrane localization was dependent on Arc higher order oligomerization since none of the oligomerization-defective truncated mutants (NTD-linker and linker-CTD) showed a similar phenotype (Fig. S4A).

Also, even if Arc NTD α-helices are predicted to interact with lipid bilayers, both through their positive net charge and palmitoylation ^18, 40, 41^, we observed that Arc truncated mutant missing the CTD (NTD-linker) diffuses in the cytosol as a tetramer and no plasma membrane localization is detectable (S1J, K). Does the interaction of Arc with the endocytic machinery rely on membrane recruitment above a certain *n*-meric state? And what drives Arc’s membrane recruitment? We performed atomistic molecular dynamic simulations to probe the strength of Arc NTD interaction with lipid membrane and to investigate whether palmitoyl groups play an important role. Since the structure of Arc NTD has been experimentally determined at low resolution only in SAXS^17^, we used the deep learning method TrRosetta^42^ to predict Arc-NTD (24 -134 aa) (Fig. S4B, upper panel). Then, four NTDs (non-palmitoylated, palmitoylated, palmitoylated with 1 or 3 carboxylic tails pre-inserted) were positioned next to a simulated lipid bilayer, which mimicked the physiological lipid composition of the inner plasma membrane leaflet (Fig. 4A). The number of residues in contact with lipids (within 3.5 Å of any of the lipids’ atoms) rapidly increased within nanoseconds after the simulation onset and stabilized between values 15-30, regardless of the NTD variant. This suggested that Arc NTD can, even without palmytoilation, strongly interact with lipids (Fig. 4B, left panel). From the simulation, phosphatidyl-ethanolamine (PE) and phosphatidylinositol 4,5-bisphosphate (PIP2) were the lipids most in contacts with the NTDs (Fig. 4B, right panel). The negatively charged PIP2 lipids formed the majority of contacts despite their considerably lower abundance (10 mol%) compared to PE lipids (40 mol%). Interestingly, despite their negative charge, phosphatidyl-serine (PS) lipids formed) lipids made fewer contacts than PIP2, despite being more abundant (15 mol%). This observation suggests that the presence of PIP2 lipids increases the interaction between membrane and NTD. Furthermore, coarse-grained molecular dynamics simulation showed that lateral diffusion of NTDs along the lipid bilayer led to the accumulation of PIP lipids, corroborating the strong interaction between NTD and PIP (Fig. S4B, lower panel).

**Fig. 4.**
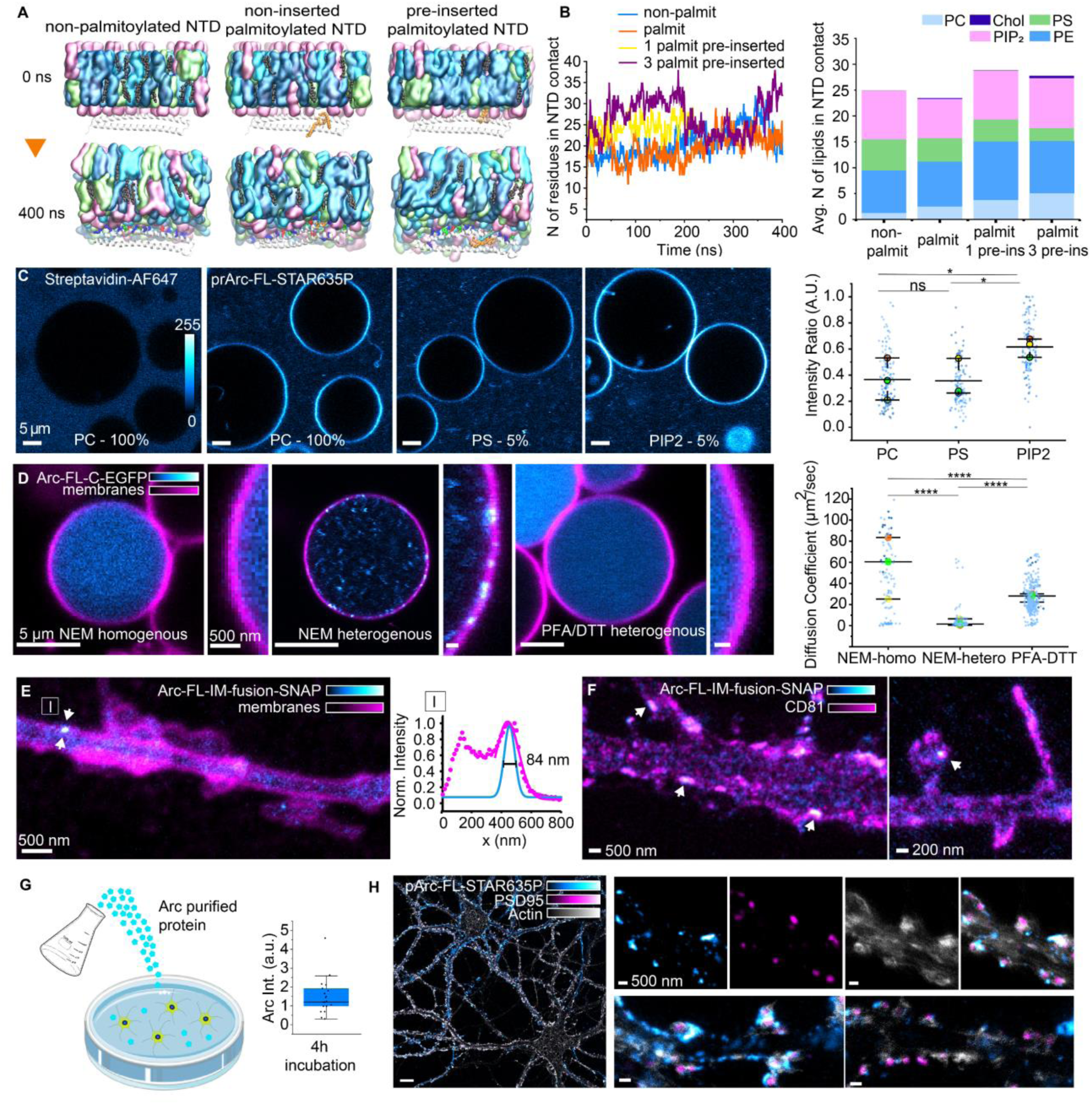
Palmitoylation and PIP lipids modulate but they are not required to mediate Arc-membrane interaction. (A) Atomistic simulation of Arc-NTD (24-134 aa)-membrane interaction. Arc-NTDs are placed in proximity to lipid bilayers composed of PE, PS, PC, PIP2 lipids and cholesterol. Arc NTDs are respectively non-palmitoylated, triple palmitoylated but not pre-inserted in lipid bilayer or triple palmitoylated and pre-inserted. Simulation duration: 400 nsec. (B) (left panel) Graph showing the number of Arc-NTD residues in contact with lipids over the time of the simulation for the different conditions. (right panel) Graphs showing the average number of lipids per type in contact with Arc-NTDs for the different conditions. (C) (left panels) GUVs incubated with streptavidin-AF647 or rat prArc-FL-STAR635P show Arc interaction with lipid bilayers for distinct GUV lipid composition (100% PC, 95% PC + 5% PS or 95% PC + 5% PIP2). (right panel) Plot showing the ratio between Arc intensity at GUVs and Arc intensity in PBS solution for the three GUV compositions. PIP2 GUVs Arc intensity ratio is significantly higher than in PC and PS conditions. The plots’ the middle line represents the mean, and the whiskers are derived from 1.5 * interquartile range. Two-sample two-sided Kolmogorov–Smirnov test p-values: PC – PS p-value 0.97, PS – PIP2 p-value 0.0326, PC – PIP2 p-value 0.0326, from 3 independent experiments whose mean values are color-coded). (D)(left panel) Representative images of HeLa cells Arc-FL-C-EGFP-derived GPMVs. Arc-FL-C-EGFP is in cyan the plasma membrane, labelled with red dye conjugated phospholipids in magenta. Two phenotypes were isolated from NEM derived GPMVs, homogeneous (N=27) and heterogeneous (N=44), for DTT-derived vesicles only homogeneous phenotype was observed (N=68). (right panel) FCS on GPMVs populations showing significantly different distributions of D_t_s (mean Dt_NEM-homogeneous:_ 41.5 um^2^/s, mean Dt_NEM-heterogeneous_: 4.8 µm^2^/s, mean Dt_PFA/DTT_: 25.6 um^2^/s, obtained from 3 independent experiments whose mean values are color-coded. The plots show the middle line representing the mean, and the whiskers are derived from 1.5 * interquartile range. Two-sample two-sided Kolmogorov–Smirnov test p-values KS test: PFA/DTT - NEM homo p-value: 2.16×10^-10^, PFA/DTT -NEM_hetero p-value 9.49×10^-64^, NEM_homo - NEM_hetero p-value 8.97×10^-21^). (E) Representative neurite from a primary neuron (DIV7) expressing Arc-FL-IM-fusion-SNAP (cyan), cell membranes are labelled using the lipophilic dye NileRed (magenta). Arc nanoclusters proximity to plasma membrane is assessed via 2-color live STED nanoscopy, an 84 nm FWHM Arc nanocluster, which colocalizes with NileRed staining, is highlighted. (F) Representative neurite of a primary cortical neuron (DIV16) expressing Arc-FL-IM-fusion-SNAP (cyan) and CD81-EGFP (magenta): a subset of Arc nanoclusters (arrows) colocalizes with CD81. (G)(left panel) Cartoon illustrating the experiment workflow of adding rat prArc-FL-STAR635P from bacteria to neuronal cultures for 4h. The incubation results in a neuron-to-neuron dependent interaction quantified by mean intensities variation and reported in the box plot (right panel). The box plot shows the 25–75% interquartile range, with the middle line representing the mean, and the whiskers derived from 1.5 * interquartile range. (H) Representative primary cortical neurons (DIV21) incubated for 4h with rat prArc-FL-STAR635P (4 ug) (cyan) and immunostained against PSD95 (magenta) and labelled for actin (gray). Arc is shown in proximity to the extracellular leaflet of neuronal plasma membranes accumulating at the level of dendritic spines (upper and low left panels), while for other less accumulation is observed (lower right panel).

The simulations were experimentally supported by *in vitro* experiments, in which Giant Unilamellar Vesicles (GUVs) were incubated with non-palmitoylated rat Arc protein purified from bacteria (prArc-FL) and subsequently labelled via STAR635P dye (Fig. 4C, left panels). Arc accumulated at the lipid bilayer of GUVs while Streptavidin-Alexa647, used a control, did not show any tendency to interact with lipids. As additional control, we incubated GUVs solely with either NHS-STAR635P dye or EGFP, and localization at GUVs membrane was not observed (Fig S4E, F). This showed that Arc-FL directly associates with lipid membranes independently from palmitoylation.

To assess experimentally the effect of negatively charged lipids over Arc-membrane association, the lipid composition of GUVs was modified (Fig. 4C, central panels). Consistently with the simulation, 5% PIP2 lipids in 95% PC, but not 5% PS, were enough to significantly increase the ratio between Arc at GUV membrane and in solution (Fig. 4C, right panel). Altogether these results showed the capability of Arc to interact with lipids and preferentially with PIP2 in an *in vitro* system.

Since Arc protein as purified from bacteria intrinsically lacks palmitoylation, and GUVs do not replicate exactly the membrane composition of a cellular environment, we took advantage of a more complex system to study the effect of palmitoylation over Arc phenotype: Giant Plasma Membrane Vesicles (GPMVs). GMPVs are vesicles derived from cellular plasma membrane which lack organelles, cytoskeletal components, and PIP lipids; hence they can be used as a clean system to study membrane and protein interaction dynamics. In addition, the palmitoylation status can be modified using distinct vesiculation chemicals: NEM (N-Ethylmaleimide) preserves palmitoylation while DTT (dithiothreitol) together with PFA (paraformaldehyde) removes the palmitoyl tails from proteins^43^. GPMVs were produced from HeLa cells expressing Arc-FL-C-EGFP. With confocal resolution, two populations of GPMVs were observed from NEM treated cells (Fig. 4D, left panel and central): GPMVs with Arc homogenously distributed and GPMVs with Arc organized in micrometric clusters and in patches in proximity to the plasma membrane. With PFA-DTT, Arc showed only a homogeneous distribution (Fig. 4D, right panel). From FCS data analysis, Arc *n*-meric states were inferred from diffusion coefficients. For NEM-treated heterogeneous population a broad distribution of states, from monomers to high-order oligomers (on average size 627 Arc molecules) was observed. For the NEM-treated homogeneous population, Arc average diffusion coefficient (D_t_ = 41.5 μm^2^/s) was comparable with the estimated diffusion coefficient of the monomeric Arc-FL-C-EGFP (D_t_ = 40.5 μm^2^/s). For DTT-treated GPMVs, instead, the measured diffusion coefficient (D_t_ = 25.6 μm^2^/s) corresponded on average to the diffusion of a tetramer (4.3 units). Additional experiments with the truncation mutant Arc linker-CTD confirmed that neither NEM nor PFA/DTT treatments induced artificial clustering (Fig. S4C). These results further demonstrated that the lack of PIP lipids in NEM-derived GPMVs did not affect the capability of Arc to interact with the membrane and oligomerize; however, in DTT-derived GPMVs, where neither PIP lipids nor palmitic anchors were present, no membrane recruitment was observed, and on the contrary an average population of low-order oligomers (tetramers) was measured.

Experiments in GPMVs demonstrated that palmitoylation is nevertheless important for Arc membrane localization. Thus, we performed additional coarse-grained molecular dynamics simulations considering palmitoylated and non-palmitoylated NTDs positioned in solution close to a PIPs-free lipid bilayer (Fig. S4D). We observed that palmitoylated NTDs interacted first with the membrane and ultimately with each other, whereas non-palmitoylated NTDs interacted with each other first and eventually with the membrane. These results suggested that, although palmitoylation is not essential for Arc-membrane interaction, it increases the probability of Arc to come in proximity to the membrane and oligomerize. Moreover, we expressed Arc-FL with mutated palmitoylation site (Arc-C94-98A) in HeLa cells where PIP lipids were present, and we observed an increase in the cytosolic homogenous phenotype at the expense of membrane interaction and oligomerization (Fig S4H). Overall, we concluded that neither the presence of palmitoyl tails nor PIP lipids is necessary for Arc-lipid interaction. However, in a cellular environment, in the absence of both palmitic anchors and PIP lipids, the cytosolic interaction between NTDs prevails over NTD-lipid interactions, thereby precluding high-order oligomerization.

To evaluate if these observations can be extended to neuronal cells, GPMVs were produced from mEGFP-N-Arc expressing neurons. Similar phenotypes to HeLa cells were observed: GPMVs with homogenous Arc, and Arc organized in micrometric clusters and in proximity to the plasma membrane (Fig. S4G).

Arc-plasma membrane localization was also assessed in neurons using 2-color live STED imaging. Neurons expressing Arc-FL-IM-fusion-SNAP had a subset of Arc nanoclusters in close proximity to the plasma membrane, labelled either with NileRed (Fig. 4E), or with the transmembrane protein and exosomes marker CD81 (Fig. 4F). Ultimately, the ability of Arc to interact with lipids in neuronal cells was further confirmed by incubating primary cortical neurons with prArc-FL-STAR635P (Fig. 4G). Upon 4h of chronic incubation, Arc was shown interacting with the extracellular leaflet of neuronal plasma membrane, it accumulated at the level of dendritic spines, which were labeled via actin and PSD95 (Fig. 4H, right higher panels). Interestingly, the interaction was evident already after 20 min post-incubation (Fig S4I) and it was highly variable from neurons to neurons (Fig. 4H, right lower panels).

Taken together the data showed that Arc, above low-order oligomers, can interact directly with membranes in cells and in neurons, the interaction is favored by the presence of negatively charged PIP lipids, and positively modulated by palmitoylation.

### Arc induces membrane inward bending and high-order oligomers affect AMPA receptors surface levels

So far, we observed different *n*-meric states of Arc in neurons: fast diffusive low-order oligomers, low-order oligomers in proximity to the PSD95 lattice, high-order oligomers sized 60-90 nm localized mostly in dendritic spines, and Arc semi-circular organizations, which populated the EZ. Arc was also interacting with lipid bilayers, and in proximity to the plasma membrane in neurons and in HeLa cells. Considering these observations and Arc homology to GAG polyprotein, we hypothesized that Arc might oligomerize at the plasma membrane level to mediate lipid bilayer bending favoring AMPA receptors internalization. Plasma membrane availability, PTMs such as palmitoylation, and Arc oligomerization above low-order oligomers should be critical in the process.

To test this hypothesis, we first performed coarse-grained simulations of Arc-NTDs-plasma membrane interaction, and we observed that Arc-NTDs show a tendency to increase local membrane curvatures (Fig. 5A). In addition, incubating prArc-FL-STAR635P with GUVs resulted in the formation of inward and outward tubulations (Fig. 5B, left panel). The tubulations reached micrometers in length, the majority were facing outward (37% among the total GUVs analyzed), a minority (9%) were facing inward and 15% of them showed both inward and outward tubulations (Fig. 5B, right panel).

**Fig. 5.**
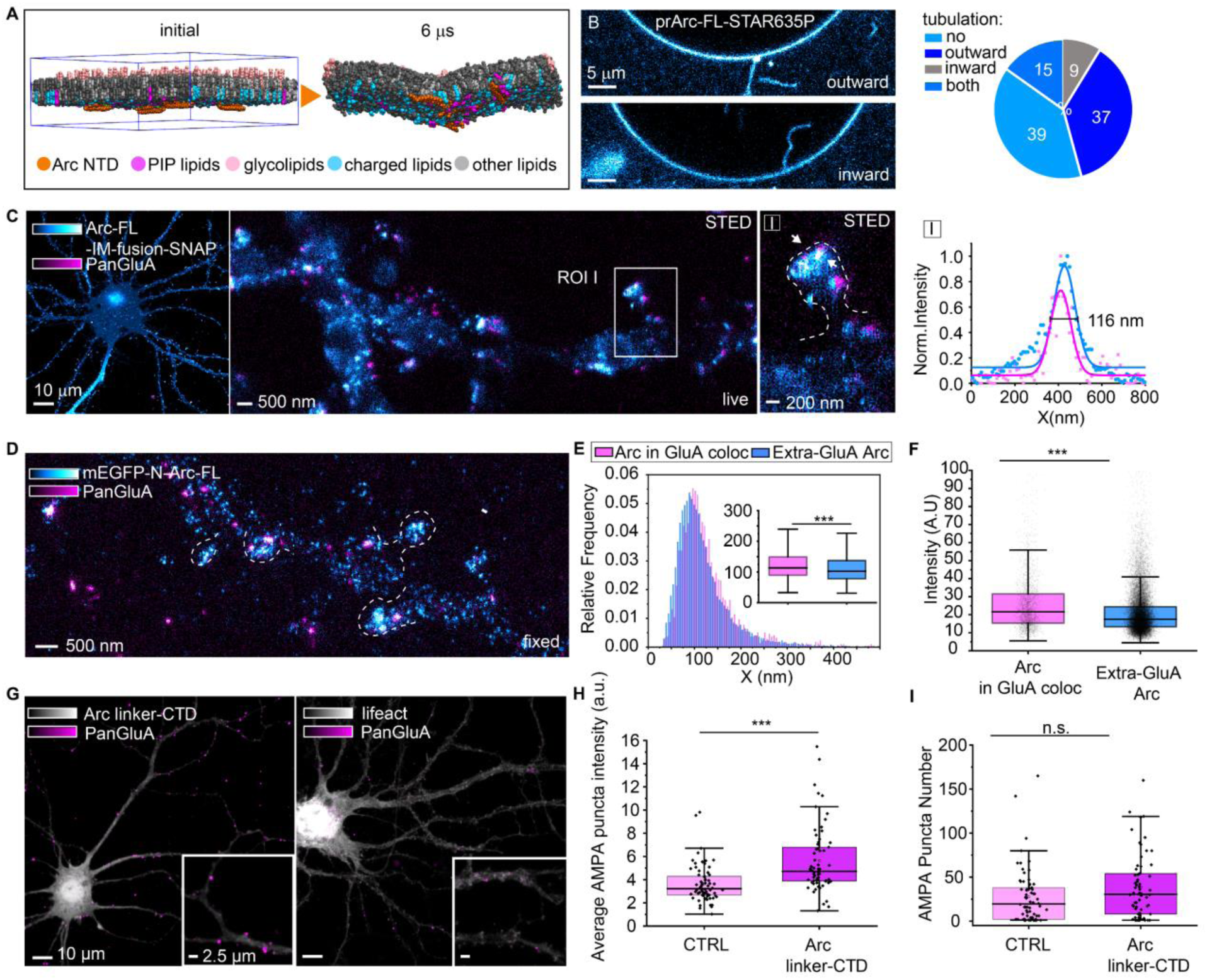
Arc induces membrane inward bending and high-order oligomers affect AMPA receptors surface levels. (A) Snapshots from a coarse-grained molecular dynamic simulation of 4 triple palmitoylated Arc-NTDs inserted into a membrane mimicking the asymmetric lipid composition of the plasma membrane. (B) (left panel) Representative images of PC GUVs incubated with prArc-FL-STAR635P showing inward and outward tubulations. (right panel) Pie chart showing inward and outward tubulation percentages. Among the analyzed GUVs, 39% (N=116) presented no tubulations, 37% (N=110) had outward tubulations, 15% (N=45) had both inward and outward tubulations while 9% (N=26) showed inward tubulations. (C) Representative primary cortical neuron (DIV20) genetically encoding Arc-FL-IM-fusion-SNAP (cyan) and live immunostained for PanGluA (magenta). In ROI I, Arc nanoclusters of different sizes localize in spines’ heads in proximity and co-localizing with the AMPA receptor. Arc nanoclusters of 116 nm FWHM is shown to co-localize with GluA, the line profile is traced along the white arrows. (D) Representative mature hippocampal neurite (DIV21) expressing mEGFP-N-Arc-FL (cyan) and PanGluA (magenta), live stained prior to fixation. (E) The histogram reports the FWHM distributions of mEGFP-Arc nanoclusters co-localizing with GluA (pink) and the extra-GluA(blue) population (24 DIV21-22 primary hippocampal neurons from 2 independent cultures, N=25080 fitted nanoclusters). As a histogram insert, the box plot shows that Arc nanoclusters in colocalization with GluA are significantly bigger than extra-GluA ones. Box plots show the 25–75% interquartile range, with the middle line representing the mean, and the whiskers derived from 1.5 * interquartile range. Two-sample two-sided Kolmogorov–Smirnov test p-value: 1.6117×10^-25^). (F) The box plot shows that Arc nanoclusters co-localizing with GluA (pink) are significantly brighter than extra-GluA ones (blue). Box plots show the 25–75% interquartile range, with the middle line representing the mean, and the whiskers derived from 1.5 * interquartile range. Two-sample two-sided Kolmogorov–Smirnov test p-values: 3.3691×10^-05^). (G) Representative primary cortical neurons (DIV20-21) expressing Arc linker-CTD-SNAP and Lifeact-YFP (gray) and live immunostained for GluA receptors (magenta) where GluA receptors puncta intensities can be compared. (H) The box plot shows that GluA puncta intensities are significantly higher for Arc linker-CTD-SNAP expressing neurons compared to Lifeact-YFP expressing (CTRL) neurons. Each data point represents the average AMPA puncta intensity per neuron (N_linker-CTD_= 66, N_CTRL_= 70, from 3 independent primary cortical cultures). Box plots show the 25–75% interquartile range, with the middle line representing the mean, and the whiskers derived from 1.5 * interquartile range. Two-sample two-sided Kolmogorov–Smirnov test p-value: 7.25×10^-8^. (I) The box plot shows that puncta number in Arc linker-CTD-SNAP expressing neurons are not significantly different from the number in Lifeact-YFP expressing neurons (CTRL) Each data point represents the average AMPA puncta intensity per neuron (N_linker-CTD_= 66, N_CTRL_= 70, from 3 independent primary cortical cultures). Box plots show the 25–75% interquartile range, with the middle line representing the mean, and the whiskers derived from 1.5 * interquartile range. Two-sample two-sided Kolmogorov–Smirnov test p-value: 0.2731.

To investigate Arc *n*-meric state in the endocytosis process, we localized Arc nanoclusters in respect to AMPA receptors, considering its direct interaction with Stargazin. Arc-FL-IM-fusion-SNAP was expressed in neurons, that were live stained for GluA and imaged with STED microscopy. Arc nanoclusters were found in colocalization with GluA (Fig 5C, ROI I).

To further characterize Arc nanoclusters, mEGFP-N-Arc-FL was expressed, and neurons were similarly live stained for GluA prior to fixation (Fig. 5D). Arc nanoclusters colocalizing with GluA were significantly brighter and larger in diameter (Fig. 5E) (co-localization median FWHMxy: 113.5 nm, residual population median FWHMxy: 102.5 nm) than the extra-GluA components (Fig. 5F). To exclude the fact that large nanoclusters could derive from the 3D spine geometry projection, we took advantage of 3D STED microscopy, which features axial resolution of about 70 nm, and it confirmed that Arc nanoclusters colocalized with AMPAR (Fig. S5A).

The observed Arc nanoclusters in co-localization with AMPA receptors might result from the accumulation of monomeric Arc at the level of its post-synaptic interaction partners. To clarify this, the truncated mutant Arc linker-CTD-SNAP, which preserves the binding site for PSDs partners, but does not form oligomers, was expressed in GluA live stained neurons (Fig. S5B). No Arc nanoclusters were found colocalizing with AMPA receptors in live 2-color STED. Moreover, imaging and spectroscopic quantification, also supported by AlphaFold interaction prediction, revealed that Arc truncated mutant linker-CTD co-expressed with Arc-FL hampers Arc-FL higher-order oligomerization and plasma membrane localization, resulting in the cytosolic diffusion as an hexamer (on average) (Fig. S5C-E). Therefore, considering linker-CTD’s dominant negative effect over Arc-FL, we took advantage of it to assess the function of Arc high-order oligomers in neurons.

In neurons expressing Arc linker-CTD-SNAP, GluA puncta were significantly brighter than in a control conditions, where neurons were expressing Lifeact-YFP (average norm. intensity CTRL: 3.064; average norm. intensity Arc linker-CTD-SNAP: 3.854) (Fig. 5G, H), but there was not a significant difference in the total number of puncta (Fig. 5I). This observation indicates that Arc linker-CTD affects AMPA receptor surface levels which supports a direct connection between Arc high-order oligomerization and AMPA receptor regulation.

Overall, we showed that Arc can induce bending and tubulations in lipid bilayers, that Arc nanoclusters are in close proximity to the membrane protein AMPA receptors, and that the dominant negative mutant, which affects Arc-Arc high-order intermolecular interaction and plasma membrane localization, increases AMPA receptors surface level.

## Discussion

In this work we provide the first direct visualization of Arc oligomers in neuronal cells. Super resolution microscopy and complementary spectroscopic techniques enabled to detect both low order and high order oligomers based on structural (size and shape) and dynamic differences (translational and rotational).

With STED and DNA-PAINT super resolution microscopy we resolved the Arc organization in nanoclusters in 2D and 3D. The images showed clusters of different sizes ranging from 6 to 300 nm, the number of observed clusters increases in density within the dendritic spines.

The presence of such clusters was further confirmed with time resolved anisotropy, which showed rotational diffusivity values typical of objects with diameters of 60-90 nm and larger than 200 nm. While the first population can represent Arc high order oligomers, the larger static population might represent Arc molecules binding larger structure like F-actin or post synaptic density proteins. Within the peri-synaptic area Arc is found in a diverse type of *n*-meric states. Arc low-order oligomers (from monomers to dimers/tetramers, 6-11 nm in diameter) located within PSD95 lattice. Arc structures of about 60-80 nm with a radial symmetry located across the whole spines, and Arc semi-circular structures located in the endocytic zone. The presence of low-order oligomers is in accordance with the findings of Arc monomers and dimers/low-order oligomers reported by Mergiya et al., 2023^25^. We hypothesized that the observed 60-80 nm structures might represent Arc higher-order oligomers made of 60 - 120 units described so far only *in vitro*. This is in accordance with the oligomeric size of 130 molecules reported by Eriksen et al. from *in vitro* quantifications, and partially with the 32-mers showed with Zhang et al.^18^. Quantitative techniques, such as qPAINT^44^ or RESI^45^, would be required to precisely quantify Arc copy numbers.

Arc requirement in AMPA receptor endocytosis has been extensively characterized, but its *n*-meric state has never been assessed. Arc oligomerization in the form of 32-mers might be needed to promote binding to its low affinity interacting partners such as Stargazin^18^. Eriksen et al. demonstrated that Arc oligomerization motif mutant was, indeed, unable to induce transferrin endocytosis in HeLa cells. In addition, Arc was shown to interact with several proteins that have a BAR domain, protein structure that senses and stabilizes membrane curvatures, such as Endophilin-3 and PICK-1^7, 8, 27^. PICK1 was displayed, using FLIM-FRET, to interact directly with Arc at the level of the plasma membrane promoting its dimerization. Moreover, Byers et al., ^15^ reported that *in vitro* oligomerization of Arc promotes Dynamin-2/3 polymerization, which favors its GTPase-dependent endocytic vesicles abscission. Here we showed that Arc above low-order oligomeric state interacts directly with lipids in cells and in neurons, in a fashion modulated by the palmitoylation, and by the presence of PIP2 (Fig. 4). The properties of Arc-membrane interaction were first assessed via atomistic simulation and further validated in GUVs. Arc interacts with neutral lipid bilayers, but the interaction is favored by PIP2 lipids. The post translational modification of palmitoylation is shown to be not required for Arc membrane interaction but, as seen from experiments in GPMVs and coarse-grained simulation, it favors membrane interaction counteracting Arc-Arc interaction in the cytosol. The palmitoylation might, therefore, be required to facilitate Arc low-order oligomers’ recruitment at the plasma membrane, which would act as a hub to promote further high-order oligomerization.

In neurons, a subset of Arc nanoclusters was shown to be near plasma membranes and the transmembrane protein and exosome marker CD81, in addition to their membrane localization in neuronal-derived GPMVs. The direct lipid bilayer affinity of Arc was also shown by the accumulation of the purified protein at the outer leaflet of neuronal plasma membranes (Fig. 4). Furthermore, we saw both in molecular dynamic simulation and in incubation with GUVs, that Arc induces outward tubulations of lipid bilayers (Fig. 5). It is important to remember that, in these experimental settings, the outer surface of the GUVs represents the inner leaflet of cell membrane. Hedde et al.,^40^ performed similar experiments, but using monomeric Arc, and saw limited Arc-lipid interactions and just inward vesiculation within the GUVs. They pointed at the budding out in extracellular vesicles release. In our case, both inward and outward tubulations were observed but the majority were outward, suggesting an involvement in endocytic cellular processes. Concurrently, we reported the localization of Arc radially symmetric structures and semi-circular organizations within the post-synaptic EZ near Clathrin, but rarely co-localizing, which suggests no direct interactions. Moreover, Arc colocalizes with AMPA receptors, while the dominant negative mutant, which affects high-order oligomerization and membrane interaction, resulted in an increase of the AMPA receptors surface level.

Based on our observations, and the Arc structural homology with GAG protein, we suggest a model in which Arc oligomerization process, similarly to HIV-1^46^, occurs in contact with the plasma membrane but causing an inward bending to favor AMPA endocytosis. We saw no Clathrin presence in proximity to Arc in the EZ. This might suggest either an early involvement of Arc in the endocytic process, or a Clathrin-independent action of Arc oligomers in AMPA endocytosis. Further experiments are required to demonstrate Arc-mediated Clathrin-independent mechanisms, but this might indicate potential involvement of Arc oligomers in alternative mechanisms of long-term depression.

In this work, we provided evidence of distinct Arc *n*-meric states and their functional role in driving membrane interaction and AMPA receptor endocytosis, but it does not exclude its release in extra cellular vesicles. In fact, we saw that Arc induces GUV tubulations both inward and outward. The inward tubulations might suggest the capability of Arc to oligomerize in a similar fashion to HIV GAG outwardly, the local plasma membrane curvature or the presence of different interacting partners may determine the directionality.

Another hypothesis, considering AMPA receptors endocytic/recycling pathway, and EVs process of release, could involve multi-vesicular bodies (MVBs). MVBs are membrane organelles which contain intraluminal vesicles (ILVs), which are formed via the inward budding of late endosomal membranes^47, 48^. MVBs content can be either degraded via fusion with lysosomes or released extracellularly as exosomes. Arc oligomers still bound to AMPA accessory subunits, facing the cytosolic side of endosomes may end up trapped into ILVs, which would be then exocytosed. Indeed, here we saw colocalization between Arc nanoclusters and the exosome marker CD81; and multiple studies demonstrated how EVs are released upon synaptic potentiating and depressing stimuli, similarly to Arc containing EVs^49–51^. On this line, it would be informative to study the protein composition of Arc-containing EVs and to further scrutinize for the presence of AMPA-associated proteins.

## Acknowledgments

We thank the Protein Science Facility at Karolinska Institute for the protein purification. Francesca Pennacchietti for the critical reading, discussion and guidance and Michael Ratz for the cloning guidance. I.T. thanks the ERC-CoG (Inspire 101002490) for supporting the research.The molecular dynamics simulations were performed on resources provided by Human Frontier Science Program (RGP0025/2022) and the Swedish Research Council Starting Grant (grant no. 2020-02682). We thank the SciLifeLab Advanced Light Microscopy facility and National Microscopy Infrastructure (VR-RFI 2016-00968) for their support on imaging. Computing at High Performance Computing Center North (HPC2N, supercomputer Kebnekaise) and the Swiss National Supercomputing Centre (supercomputer Piz Daint).

## Authors contributions

I.T. conceived research and supervised the project. M.D. cultured and labeled neurons, optimized the labeling protocols, performed the cloning and the imaging experiments. G.C. cultured the neurons, produced the viruses and optimized the labelling and immunostaining protocols. M.S. cultured the neurons, the glial cells, optimized the labeling protocols and the Calcium-Phosphate transfections protocol. J.A. built the STED optical set-up and performed the Arc nanoclusters density analysis. C.S. performed the imaging experiments with EVs markers together with M.D. E.S. produced and performed the GUVs, GPMVs and FCS experiments together with M.D. L.R performed the molecular dynamics simulations supervised by L.D. L.A.M, E.M.U. and R.K. performed DNA-PAINT experiments and data analysis together with M.D. supervised by R.J, A.V. performed the STARSS measurement together with M.D. M.D. analyzed the data and wrote the manuscript together with I.T. and with the input from all the authors.

## METHODS

### RESOURCE AVAILABILITY

#### Lead contact

Further information and requests for resources and reagents should be directed to and will be fulfilled by the lead contact, Ilaria Testa.

### Material Availability

Plasmids generated in this study will be deposited to Addgene and reported with the name and catalog number or unique identifier.

### Data and Code Availability

- Microscopy data reported in this paper will be shared by the lead contact upon request.
- All original code is available in this paper’s supplemental information.
- Any additional information required to reanalyze the data reported in this paper is available from the lead contact upon request.

## EXPERIMENTAL MODEL AND SUBJECT DETAILS

All experiments were performed in accordance with animal welfare guidelines set forth by Karolinska Institutet and were approved by the Swedish board of agriculture (Jordburks verket). Rats were housed with food and water available *ad libitum* in a 12-hour light/dark environment.

### Primary mature neuronal culture

Primary cultures were prepared from embryonic day 18 (E18) Sprague Dawley rat embryos. The pregnant mothers were sacrificed with CO_2_ inhalation and aorta cut, and brains were extracted from the embryos. Cortexes or hippocampi were dissected and mechanically dissociated in Minimum Essential Medium, MEM (Thermo Fisher Scientific, 21090022). 2 x 10^5^ cells per 60 mm culture dish were seeded on poly-D-ornithine (Sigma Aldrich, P8638) coated #1.5 18 mm glass coverslips (Marienfeld, 0117580), and were let to attach in MEM with 10% horse serum (Thermo Fisher Scientific, 26050088), 2 mM L-Glut (Thermo Fisher Scientific, 25030-024) and 1 mM sodium pyruvate (Thermo Fisher Scientific, 11360-070), at 37°C at an approximate humidity of 95–98% with 5% CO_2_. After 2–4 h, coverslips were flipped over an astroglial feeder layer (grown in MEM supplemented with 10% horse serum, 0.6% glucose, and 1% penicillin-streptomycin) and maintained in Neurobasal (Thermo Fisher Scientific, 21103-049) supplemented with 2% B-27 (Thermo Fisher Scientific, 17504-044), 2 mM L-glutamine and 1% penicillin– streptomycin. The neuronal cultures were treated with 5 μM 5-fluorodeoxyuridine (FDU) at DIV 2–3, to prevent glia overgrowth. The cultures were kept for up to 24 days and fed twice a week by replacing one-third of the medium per well: up to DIV7 with Neurobasal complete medium, and after DIV7 with BrainPhys (STEMCELL tech. 05790), 1% Pen/Strep (Gibco 15140-114) and SM1 Supplement (STEMCELL tech. 05711). For experiments performed with immature cortical neurons (DIV 6-9), 100 x 10^3^ cells per well were seeded in 12 well plates on a poly-D-ornithine coated #1.5 18 mm glass coverslips. 3 hours post-plating the media was changed to Neurobasal Medium supplemented with 2% B-27 (Thermo Fisher Scientific, 17504-044), 2 mM l-Glutamine and 1% Penicillin-Streptomycin (Sigma Aldrich, P4333).

### Cell lines culture

HeLa (ATCC CCL-2) cells were cultured in DMEM (Thermo Fisher Scientific, no. 41966029) supplemented with 10% (vol/vol) fetal bovine serum (Thermo Fisher Scientific, no. 10270106), 1% penicillin/streptomycin (Sigma-Aldrich, no. P4333) and maintained at 37 °C and 5% CO_2_ in a humidified incubator. Cells were plated on #1.5 18 mm glass coverslips (Marienfeld, 0117580) 48h before imaging.

## METHOD DETAILS

### Plasmid constructs

Plasmids used for Arc exogenous expression in immature neurons and HeLa cells are: EF1a_Arc-FL-IRES-WGA-Cre, pAAV-EF1a_Arc-FL-C-SNAP-IRES-WGA-Cre, pAAV-EF1a_Arc-IM-fusion-SNAP-IRES-WGA-Cre, pSFV-SCA_Arc-C-AlfaTag, pAAV-EF1a_Arc-NTD-linker-SNAP-IRES-WGA-Cre, pAAV-EF1a_Arc-linker-CTD-SNAP-IRES-WGA-Cre, pAAV-EF1a_Arc-C-rsEGFP2-IRES-WGA-Cre and pCAG-Arc-C-EGFP, pAAV-EF1a_ArcC94-98S-C-SNAP-IRES-WGA-Cre, pAAV-EF1a_ArcC94-98S-C-SNAP-IRES-WGA-Cre. Plasmids used for Arc exogenous expression in mature neurons are pSFV-SCA_5’-UTR_Arc-IM-fusion-SNAP_3’UTR, pSFV-SCA_5’-UTR_mEGFP-Arc_3’UTR, and pGL4.11-Arc7000-mEGFP-Arc-UTRs (Kawashima et al., 2009, kindly provided by Prof. H. Bito from Department of Neurochemistry, Graduate School of Medicine, University of Tokyo). pcDNA3.1_mm_Cd81-EGFP for neuronal expression was a gift from Michael Ratz (Karolinska Institute). Plasmids were prepared from the transformants and verified via Sanger sequencing. Details of the plasmids cloning are provided in Supplementary Note 1.

### Neuron transfection and live cell staining

DIV6-9 neurons were transfected using Lipofectamine 2000 Transfection Reagent (Thermo Fisher Scientific, 11668019), according to the instructions of the manufacturer. For labelling fusion proteins, 24 hours after transfection, neurons were washed in Artificial CerebroSpinal Fluid (ACSF) and labelled with 5 μM of the SNAP substrates (New England BioLabs, SNAP-Cell 647-SiR), for 45 minutes at 37°C. Then, neurons were washed three times with ACSF and put back in the conditioned medium for at least 20 minutes. Mature primary cortical neurons were infected with a modified Semliki Forest Virus 16-18 hours before the experiment. Alternatively, neurons were transfected using calcium phosphate co-precipitation protocol as reported^52^. In short: DNA (2 μg per coverslip) was diluted in TE solution (Tris-HCl, pH 7.5, 10 mM; EDTA, pH 8.0, 1 mM). CaCl_2_ (2.5 M in 10 mM HEPES) was added to a final concentration of 250 mM. The mixed solution was added to 2× HEBS (HEPES Buffered Saline, pH 7.2). Neurons were pre-incubated in 200 μl of conditioned medium from their culture dish with 50 μl of 5× Kynurenic acid stock (10 mM dissolved in unsupplemented culture medium) in a well of sterile MW12 and placed back in the incubator until the precipitate was ready. The precipitate was then added dropwise to the cells and incubated for 3–4 h. In order to stop the transfection, a 5:1 mix of BrainPhys without SM1 and Kynurenic acid was pre-warmed. Then, 5 M HCl was added until the solution turned yellow. After removal of the transfection medium, the acidic medium was added to each coverslip which was further incubated at 37 °C/5% CO_2_ for 15–20 min. After the incubation period the neurons were transferred back to the original Petri dish containing the conditioned medium and the construct was let express for 24–48 h at 37 °C/5% CO_2_.

Imaging was performed in ACSF at RT. prArc-STAR635P was incubated (4 ug/ml) from 20 min to 4h prior to fixation.

### HeLa transfections

For transfection, 2 × 10^5^ cells per well were seeded on coverslips in a six-well plate. After one day cells were transfected using FuGENE (Promega, E2311) according to the manufacturer’s instructions. 24–36 h after transfection cells were washed in phosphate-buffered saline (PBS) solution, placed with phenol-red free Leibovitz’s L-15 Medium (Thermo Fisher Scientific, 21083027) in a chamber and imaged.

### GPMVs production

To produce GPMVs, HeLa cells were plated in 60 mm petri dishes and transfected to express Arc-C-EGFP. Cells were washed twice with GPMVs buffer medium (150 mM NaCl, 10 mM Hepes, 2 mM CaCl). Cells in GPMVs buffer either adding 2 NEM (1M) to reach 2 mM final NEM concentration or 4% PFA and DTT (1M) to reach final concentration respectively of 25 mM for PFA and 2 mM for DTT. Cells were put back at 37°C at an approximate humidity of 95–98% with 5% CO_2_ for at least 1h. GPMVs production visually assessed in a tabletop transmission light microscope. Arc-C-EGFP expressing HeLa cells-derived NEM GPMVs further subdivided in two populations: homogeneous population, in which there were no resolvable Arc-C-EGFP clusters; and heterogenous population where instead, we could resolve Arc-C-EGFP clusters.

### Rat Arc protein purification

The construct psfARC-c001 (pGEX-6p1-GST-ArcFL, plasmid #119877 obtained from Addgene) was transformed into *E. coli* BL21 (DE3) T1R pRARE2 cells. The cells were cultivated in 3000 ml Terrific Broth (TB) medium supplemented with 8 g/l glycerol and Ampicillin (50 μg/ml), Chloramphenicol (34 μg/ml). Inoculation of overnight cultures from fresh transformants followed the cultures grown at 30°C, 175 RPM overnight in the presence of 0.4% glucose. After 24h the cultures were grown in the LEX system. At different times, the OD was measured for the cultures and the temperature was set to 18°C at OD 2. The protein expression was induced at approximately OD 3 (IPTG, final concentration 0.5 mM). Protein expression continued overnight before the cells were harvested by centrifugation (10 min at 4500 × g). The lysis buffer (100 mM HEPES, 500 mM NaCl, 10 mM imidazole, 10% glycerol, 0.5 mM TCEP, pH 8.0 (1.5 ml buffer per gram cell pellet) and the complete stock solution (1 ml per 1.5 l culture: 1 tablet Complete EDTA-free (protease inhibitor cocktail, Roche) and 50 μl benzonase nuclease cell resuspension (PSF) per 1 ml) were added and the cell pellets were re-suspended on a shaker table (cold room). The resuspended cell pellets were frozen at -80°C. The frozen cell pellets were briefly thawed in water (room temperature) and cells were disrupted by pulsed sonication (4s/4s, 4 min, 80% amplitude). The sonicated lysates were centrifuged (20 min at 49000 × g) and the soluble fractions were decanted and filtered through 0.45 μm filters. The clarified lysate was loaded onto a GSTrap 4B 5 ml column (1 ml/min, GE Healthcare) and the flow-through was collected and re-loaded an additional time. The column was washed with GST wash buffer 1 (PBS, 0.5 mM TCEP, pH 7.4 and 2 (PBS, 1 M NaCl, 0.5 mM TCEP, pH 7.4, 4 ml/min). The column was then equilibrated with 10 column volumes of 3C protease cleavage buffer (50 mM Tris, 150 mM NaCl, 0.5 mM TCEP, 0.5 mM DTT, pH 8.0) (at 8°C), 3C protease (PSF, psfNTx3C-c001, 1:500 molar ratio) was added and the column was incubated overnight at 8°C. The GSTrap-column was then coupled to a 1 ml Histrap HP column (GE Healthcare), for the purpose of removing the 3C protease, and the target protein was eluted with 3C protease cleavage buffer (reverse GST purification). Finally, an elution step with GST-elution (50 mM Tris, 150 mM NaCl, 30 mM reduced glutathione, pH 8.0 (at 8°C) buffer was performed, to check if some uncleaved protein remained bound to the column. Selected fractions were examined by SDS-PAGE and fractions containing the cleaved target protein were pooled. The protein containing fractions from the reverse GSTrap purification were loaded onto the ÄKTA Xpress and purified overnight in gel filtration column HiLoad 16/60 Superdex 200 (GE Healthcare). Fractions containing the target proteins were pooled and concentrated with Vivaspin concentration filters (Vivascience 10 kDa cut off). The final concentration was measured (Nanodrop), the protein was flash frozen in aliquots of 100 or 200 μL in liquid nitrogen and stored at -80 °C. The final batch buffer was constituted by 50 mM Tris, 150 mM NaCl, pH 7.4 (at room temperature).

### NHS labelling

Arc purified protein was thawed on ice and storage buffer was exchanged to PBS in Pierce Zeba™ Desalt Spin Columns (0.5 mL, ThermoFisher). Abberior® STAR RED NHS was dissolved in DMSO at 10 mg/ml. 0.1 ml of 1M NaHCO3 (in water pH 8.5) solution was added for each 1 ml of protein (100 ug). 1 ul of the dye solution was added to the protein solution with NaHCO3 and incubated for 1 hour at room temperature with continuous stirring. Unbound dye was further removed via centrifugation in the desalting columns and the final concentration assessed via Nanodrop.

### GUV production

GUVs were produced as previously reported in Erdinc et al., 2015. Briefly, 1 mg mL^−1^ lipid solutions were prepared in chloroform (100% POPC, 95% POPC + 5% POPS, 95% POPC + 5% PIP2). Then, 5 μL of this solution were dried onto two parallel platinum wires mounted in a GUV Teflon chamber. A 300 mM sucrose solution was added to the chamber and a 10 Hz current was applied to the wires for an hour. GUV preparation was formed at room temperature. Arc-STAR635P (4 ug) was incubated with GUVs for 20 min before imaging.

### Cell membranes labelling

Cell membranes in primary neurons were labelled incubating with NileRed dye (100 mM in DMSO, Sigma-Aldrich, cat. N.72485) (1:5000) directly in the ACSF imaging. Cell membranes in HeLa cells were labelled incubating with CellBrite Red (Biotium, Cat.N 30023) at 1:200 for 20 min at 37C. GPMVs membranes were labelled for 5 min at 37C with red-PE before imaging.

### 1,6-Hexanediol (1,6-HD) treatment

Arc-C-EGFP expressing HeLa cells were treated for 1 min with different concentrations (3%, 5% and 10% w/v) of 1,6-HD (Sigma-Aldrich Cat. N 24011) in Leibovitz’s L-15 Medium (Gibco, Cat.N 21083027) supplemented with 10% FBS (Sigma-Aldrich, Cat.N. F7524) and then fixed in 4% PFA for 20 min, washed 3x with PBS and further imaged.

### Confocal Microscopy Confocal and FCS setup

Confocal imaging and FCS measurement were performed at Confocal Microscope Zeiss LSM 780 Argon laser with 488 nm lines was used to excite EGFP samples and HeNe 633 nm laser was used to excite 647SiR or JF646.

### Fluorescence Recovery After Photobleaching (FRAP) acquisition

FRAP experiments were performed in Arc-C-EGFP expressing HeLa cells using a Confocal Microscope Zeiss LSM 780 with a 63X oil immersion objective. Arc-C-EGFP micrometric clusters identified in the cytosol were either fully bleached or bleached internally but for region of interests (ROIs) with constant diameter. In both cases bleaching was carried out using Argon laser with 488 nm line at 50% of its maximum and 405 nm line at maximum power. Five images were acquired before bleaching and the recovery was monitored for 30 sec. The measurements were performed at 37C.

### STED microscopy Microscope setup for STED

STED nanoscopy has been performed at the Leica TCS SP8 3X STED, which is equipped with a HC PL APO 100x/1.40 Oil STED White objective or at a custom-built STED setup, based on a STED setup previously described^53^. *STED:* Excitation of the red-shifted dyes was done with a pulsed diode laser at 640 nm with a pulse width of 60 ps (LDH-D-C-640, PicoQuant, Berlin, Germany), excitation of the red dyes was done with a pulsed diode laser at 561 nm with a pulse width of 60 ps (PDL561 Abberior Instruments). Depletion was done with a pulsed 775 nm laser beam with a pulse width of 530 ps (KATANA 08 HP, OneFive GmbH, Regensdorf, Switzerland). Fast on and off control of the excitation and depletion lasers for STED is done using an AOTF (AOTFnC-400.650-TN + MPDS4C-B66-22-74.156, AA Opto Electronic, Orsay, France) and AOM (MT110-B50A1.5-IR-Hk + MDS1C-B65-34-85.135-RS, AA Opto Electronic, Orsay, France) respectively. The depletion beam is shaped using a vortex phase mask on a spatial light modulator (LCOM-SLM ×10468-02, Hamamatsu Photonics, Hamamatsu, Japan). A λ/4 and a λ/2 wave plate are used to create the circular polarization necessary for optimal depletion focus formation. Fast galvanometer mirrors are used for scanning in STED imaging (galvanometer mirrors 6215H + servo driver 71215HHJ 671, Cambridge Technology, Bedford, MA, USA) in a scanning system allowing constant resolution across an 80 x 80 μm^2^ FOV as described previously^53^. The fluorescence is decoupled with a dichroic mirror into two channels. Channel 1 (magenta) has a notch filter (NF03-785E-25, Semrock), a bandpass filter (ET705/100 m, Chroma), and the fluorescence is focused onto a free space APD (SPCM-AQRH-13-TR, Excelitas Technologies, Waltham, MA, USA). Channel 2 (green) has a notch filter (ZET785NF, Chroma), a bandpass filter (ET615/30 m, Chroma), and the fluorescence is focused onto a 62.5 *μ*m core diameter multi-mode fibre (M31L01, Thorlabs) coupled to an APD (SPCM-AQRH-14-FC, PerkinElmer, Waltham, MA, USA). *General:* The setup uses a 100x/1.4 oil immersion objective (HC PL APO 100x/1.40 Oil STED White, 15506378, Leica Microsystems, Wetzlar, Germany) and a microscope stand (DMi8, Leica Microsystems). The system also uses a mechanical stage for moving of the sample in the lateral dimensions (SCAN IM 130 x 85 – 2 mm, Märzhäuser, Wetzlar, Germany) and a piezo to move the sample along the axial dimension (LT-Z-100, Piezoconcept, Lyon, France). The images were recorded by exciting AbberiorSTAR580 and 647SiR or AbberriorSTAR635P. Two-color STED images were recorded line-by-line, adding up 4 lines with 0.03 msec pixel dwell time. The pixel size of the images is 19.53 nm.

### Neuron Sample preparation for STED imaging

For the live labelling of AMPA receptors, 1 μl of GluA antibody (1 mg/ml, GluA antibody - 182 411, Synaptic System) and 1 μl of FluoTag-X2 anti-mouse KLC (5 µM, cat. N1202) conjugated to Abberrior STAR580 dye were pre-incubated with 98 μl of pre-conditioned neuronal medium. Neurons were, then, incubated with the AMPA labelling solution for 10-20 min in a humidified chamber at 37°C. After the incubation time neurons were left to recover for 5 min in their original medium and washed twice with artificial cerebrospinal fluid (ACSF) before the imaging or before the fixation. Before fixation cells were washed once with 1X PBS, pre-warmed at 37°C and fixed in pre-warm 4% PFA at room temperature for 10 minutes. After being washed twice with PBS, cells were permeabilize by incubation with 0.5% TRITON™ ×100 in PBS for 5 minutes or 0.1% TRITON™ ×100 in PBS for 10 minutes. Blocking was performed by incubation with 5% bovine serum albumin in PBS (5% BSA/PBS) for 30 minutes at RT. Primary antibodies as Anti-PSD95 antibody (Santa Cruz, sc-32290, 1:50) and Anti-Arc (Synaptic System, 156 003, 1:1000) was incubated in 5% BSA/PBS for 1 hour at RT. After three 5 min washes in PBS, the sample was incubated with donkey anti-mouse Alexa594 secondary antibody (ThermoFisher A-21203, 2 mg/ml, 1:200), goat anti-rabbit Abberior STAR635P, or Anti-GFP nanobody (FluoTag®-X4 anti-GFP conjugated with Abberrior635P, N0304) and anti-PSD95 nanobody (Nanotag, N3702-Ab580-L, 1:500) in 5% BSA/PBS for 1 hour at RT. The samples were washed in PBS for 15 min and the coverslips were mounted I Mowiol.

## DNA-PAINT

### Microscope setup for DNA-PAINT

Imaging was carried out using an inverted microscope (Nikon Instruments, Eclipse Ti2) equipped with a Perfect Focus System using the objective-type TIRF configuration with an oil-immersion objective (Nikon Instruments, Apo SR TIRF 100X, NA 1.49, oil). A 561 nm laser (MPB Communications, 1 W) was used for excitation and was coupled into a single-mode fiber and into a Nikon manual TIRF module. The laser beam was passed through a cleanup filter (Chroma Technology, ZET561/10) and coupled into the microscope objective using a beam splitter (Chroma Technology, ZT561rdc). Fluorescence light was spectrally filtered with an emission filter (Chroma Technology, ET600/50m) and imaged with an sCMOS camera (Andor, Zyla 4.2 plus) without further magnification, resulting in an effective pixel size of 130 nm after 2×2 binning. The camera readout sensitivity was set to 16-bit and the readout bandwidth to 200 MHz. The central 1024×1024 pixels (512×512 after binning) of the camera were used as the region of interest. 3D imaging was performed using an astigmatism lens (Nikon Instruments, N-STORM) in the detection path^54^. Image acquisition and microscope control was performed using µManager^55^ (Version 2.0.1). The illumination angle was set into a highly inclined illumination (HILO) mode.

### Neuron sample preparation and labeling scheme for DNA-PAINT

After fixation, neurons were quenched using 100 mM NH_4_Cl (Merck, 12125-02-9) in PBS. Then, samples were washed three times with PBS and incubated with 0.1% Triton X-100 in PBS for 20 min for permeabilization. Afterwards, the samples were washed with PBS, and 3% BSA in PBS, supplemented with 0.05 mg/ml sheared salmon sperm DNA was applied for blocking. For drift-correction gold nanoparticles (1:3 dilution in PBS) were incubated for 5 min. Primary antibodies (ClathrinHC) were incubated with secondary nanobodies (Rabbit IgG Nanotag) and direct nanobodies (GFP 1H1 and GFP 1B2 and PSD95 from Nanotag) in 300 µl antibody incubation buffer overnight onto the sample. Following the overnight incubation, the sample was washed 4 times with PBS and once with Buffer C+.

### Buffers

The following buffers were used for sample preparation and imaging:

- Buffer C+: 1× PBS, 500 mM NaCl and 0.05% Tween-20, 1x PCA, 1x PCD, 1x Trolox
- Buffer C: 1× PBS, 500 mM NaCl
- Antibody Incubation buffer: 1×PBS, 1 mM EDTA, 0.02% Tween-20, 0.05% NaN_3_, 2% BSA and 0.05 mg/ml sheared salmon sperm DNA

### PCA, PCD and Trolox

Trolox (100×) was made by the addition of 100 mg of Trolox to 430 μl of 100% methanol and 345 μl of 1 M NaOH in 3.2 ml of water. PCA (40×) was made by mixing 154 mg of PCA in 10 ml of water and NaOH and adjustment of pH to 9.0. PCD (100×) was made by the addition of 9.3 mg of PCD to 13.3 ml of buffer (100 mM Tris-HCl pH 8.0, 50 mM KCl, 1 mM EDTA, 50% glycerol).

### Nanobody-DNA conjugation via single cysteine

Nanobodies against GFP (cat: N0305), tagFP (cat: N0501), rabbit and mouse IgG (cat: N2405 & N2005) were purchased from NanoTtag Biotechnologies with a single ectopic cysteine at the C-terminus for site-specific and quantitative conjugation. The conjugation to DNA-PAINT docking sites was performed as described previously^56^. First, buffer was exchanged to 1× PBS + 5 mM EDTA, pH 7.0 using Amicon centrifugal filters (10k MWCO) and free cysteines were reacted with 20-fold molar excess of bifunctional maleimide-DBCO linker (Sigma Aldrich, cat: 760668) for 2-3 hours on ice. Unreacted linker was removed by buffer exchange to PBS using Amicon centrifugal filters. Azide-functionalized DNA was added with 3-5 molar excess to the DBCO-nanobody and reacted overnight at 4°C. Unconjugated nanobody and free azide-DNA was removed by anion exchange using an ÄKTA Pure liquid chromatography system equipped with a Resource Q 1 ml column. Nanobody-DNA concentration was adjusted to 5 µM (in 1xPBS, 50% glycerol, 0.05% NaN3) and stored at -20°C.

### Neuron imaging for DNA-PAINT

Imaging was performed in sequential rounds of DNA-PAINT with the parameters summarized in Supplementary Table 2.

## STARSS

### Microscope setup for STARSS

The STARSS experiments^23^ were performed in a custom confocal microscopy system equipped with a dual channel polarized detection (parallel and perpendicular polarizations).

### STARSS measurement

The anisotropy decay in the μ-second time domain was measured using rigid rsEGFP2 fusion of the Arc protein. At every scanning position a polarized pulse scheme of light was delivered to the sample: at the beginning of the sequence the fluorescent protein was fully switched off using 1.5 millisecond of 488 nanometer light at roughly 20 kW/cm^2^, then roughly 10% of the protein was switched on using a short burst of 250 nanosecond of linearly polarized 405 nm light at roughly 110 kW/cm^2^ (parallel direction), finally fluorescence was induced using 1.5 millisecond of circularly polarized 488 nanometer light at roughly 20 kW/cm^2^. The fluorescence photons were recorded with a binning time of 1 microsecond.

## RNA-FISH

### Probe design and production

Arc RNA FISH probes were designed based on rat transcriptome as previously reported by Gelali^57^, and they are reported in Supplementary table 1.

### iFISH

RNA FISH was performed as reported in Gelali et al., 2019. Briefly, neurons were fixed fix in 4% PFA/0.4xPBS (-Ca, -Mg), for 10 min at RT and then incubate in 1xPBS/0.5% Triton X-100 / RVC for 20 min. Cells were further incubated in in the RNA WASH 25% buffer for 5 min followed by primary hybridization mix (1 ul of primary probe, 25 uM, for 100 ul o RNA hyb buffer) for 24 h at 30 C. Wash in RNA WASH 25% buffer followed for 30 min at 30C. Same was performed for the secondary hybridization mix (1 ul of secondary probe conjugated to Alexa 594 dye, 2 uM, for 100 ul o RNA hyb buffer) with additional 60 min incubation with RNA WASH 25% buffer and a rinse with 2xSSC buffer. A standard immunofluorescence staining as described above followed to label Arc-EGFP. <u>Hybridization buffer</u> (RNA HYB 25%): 10% Dextran sulfate (Sigma #D8906), 25% Formamide (Ambion #AM9342), 2x SSC (Ambion # AM9765), 1 mg/ml E.coli tRNA (Sigma #R4251), 0.02% BSA (Ambion #AM2616), 2 mM Ribonucleoside vanadyl complex (NEB S1402S), up to 10 ml Nuclease-free water (Ambion #AM9932). <u>Wash buffer</u> (RNA WASH 25%)(pH:7-8): 25% Formamide (Ambion #AM9342), 2x SSC (Ambion # AM9765), 32.5 ml Nuclease-free water (Ambion #AM9932).

### Molecular Modeling

The prediction of the structure of Arc NTD was determined using the Robetta webserver. We employed atomistic and coarse-grained molecular dynamics simulations as well as the HADDOCK docking method. The 24-134 aa of Uniprot sequence Q63053 was predicted by the TrRosetta^58^ method available through (https://robetta.bakerlab.org) at the confidence level of 0.71. The two helices of both structures are well aligned with RMSD of 3.36 Å (CE alignment) or 2.72 Å (SALIGN), as computed in PyMod plugin in PyMol.

Both atomistic and coarse-grained molecular dynamics simulations were employed and carried out using Gromacs 2020.3 (atomistic simulations) or 2019.4 (coarse-grained)^59^. The atomistic simulations considered the NTD, either non-palmitoylated or containing palmitoyl groups at residues C94, C96, and C98, in proximity to a symmetric lipid bilayer with area 10×10 nm^2^ mimicking the composition of the intracellular plasma membrane leaflet (5% DOPC, 5% POPC, 20% DOPE, 20% POPE, 7% DOPS, 8% POPS, 10% POPI24, 25% cholesterol)^60^ bathed in a 150 mM KCl solution. All systems were built with the charmm-gui web server using the Charmm36m force-field^61^. The simulations were run in the NPT ensemble following the charmm-gui default molecular dynamics parameters for the equilibration and production steps. The production step was 400 ns long. The coarse-grained simulations were based on the MARTINI force-field version 2.6 that includes the parameters for palmitoylated cysteines^62^. Three systems were considered. In the first, 4 triple-palmitoylated Arc NTDs were positioned at the interface of a 40 × 40 nm^2^ membrane bathed in 150 mM NaCl (see Fig. 6A). The membrane consisted of an asymmetric distribution of ∼60 different types of lipids, including PIP lipids, and mimicked the realistic composition of an average mammalian plasma membrane ^63^. In the second and third system, respectively, 4 non-palmitoylated or 4 palmitoylated Arc NTDs were positioned in solution next to a 30 × 30 nm^2^ membrane devoid of PIP lipids (see Suppl. Fig. 5D). Following equilibration steps these systems were simulated for 5-10 µs in two replicas. All systems were prepared and simulated following protocols from http://cgmartini.nl/^64, 65^. No restraints were imposed on the protein secondary structure. The simulated trajectories were visualized in VMD^66^ and analyzed using custom scripts based on MDAnalysis ^67^.

Arc-FL, Arc truncated mutants and multimers structures were predicted taking advantage of Colab notebook (https://colab.research.google.com/github/deepmind/alphafold/blob/main/notebooks/AlphaFold.i <u>pynb</u>), it uses a slightly simplified version of AlphaFold v2.1.0^68^. Structures and prediction errors were then visualized using ChimeraX (v1.4rc202205290614).

## QUANTIFICATION AND STATISTICAL ANALYSIS

### Fluorescence Recovery After Photobleaching (FRAP) Analysis

Each data point in the fluorescent recovery curves the mean intensity values in the ROI for each acquired frame. The values were then normalized from 0 to 1 and fitted with a single exponential function: y = y0 + A1*(1 - exp(-x/t1)) using OriginLab 2021 software, where t1 represents the fluorescence recovery constant, A1 is the mobile fraction (M_f_) and y0 is the y-intercept. The immobile fraction (IM_f_) was calculated as follows: IM_f_ = 1-M_f_. The half time recovery (τ_1/2_) was calculated as it follows: τ_1/2_=t1*ln(2).

### Confocal Image Analysis: Arc micrometric cluster analysis

Arc micrometric clusters area and roundness were calculated with ImageJ from Arc-C-EGFP expressing HeLa cells. Gaussian Blurring (sigma: 0.5) was first applied to each image which was then further thresholded (100, 255), binarized and the ImageJ plugin Analyze Particles was applied.

### Confocal Image Analysis: AMPA puncta intensity analysis

Stacks of confocal images (separated of 250 nm) were taken from fixed samples expressing Arc-ΔNTD-SNAP or lifeact-YFP and live stained for AMPA receptors. 30×30 um manually cropped Arc or lifeact images were binarized upon thresholding using the Haung algorithm to identify neurites. Binary masks were then enlarged by 0.25 um and only AMPA signals within that area were taken into account. AMPA puncta were identifies using Maxima plugin of ImageJ (prominence: 2). The photon-count of the local Maxima defines the AMPA intensity.

### FCS Data Analysis

FCS correlation curves obtained from the cytosol of HeLa cells expressing the different mutant combinations tagged via SNAP with the ligand coupled to JF646 or Arc-C-EGFP expressing HeLa cells derived GPMVs were analysed using the software FoCuS_point (version 1.13.156). The data were fitted with a mono exponential diffusion equation (A1 fixed to 1 and AR1 fixed to 6). Diffusion coefficient (D) was then derived from the diffusion time (τ_D_) using the formula D =ω^2^/(8*lg2* τ_D_), where ω is the confocal spot diameter measured for each experimental session. Triplets were excluded moving the residual tau threshold to 10^-2^ msec.

### Arc oligomers size retrieving model from diffusion coefficients

As reported by Galiani al., 2022^69^ the transit diffusion time for a cytosolic 3x monomeric EGFP (80.85 kdal) in GPMVs is 39 ± 13.5 µm^2^/ s. We applied the Stokes-Einstein equation adapted for the masses as reported by Young et al., 1980 as it follows:

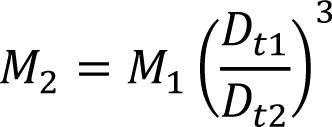

Where M_1_ and M_2_ are the respective species’ masses, while 𝐷_𝑡1_ and 𝐷_𝑡2_ their measured diffusion coefficients. From that we estimated first the diffusion coefficient of Arc-C-EGFP (72.4 kdal) monomer as being 40.5 µm^2^/ s and then the Arc *n*-meric states for each GPMVs conditions.

Similarly as above, to extract the *n*-meric states of Arc cytosolic diffusive components in HeLa cells, we first assumed Arc linker-CTD as a monomer, then dividing D_t linker-CTD_ by the D_t_ of the other species we extracted the distinct oligomeric states applying the Stokes-Einstein equation adapted for the masses.

### STED Image Analysis: Arc nanoclusters size, intensity, and marker colocalization

The analysis of Arc nanoclusters size and intensity for immature and mature primary neurons for the different Arc labelling strategies was performed using the Fiji distribution of ImageJ and using the script provided, see Code Availability. Each nanocluster was localized using the Find Maxima plugin of ImageJ (prominence: 10). The photon-count of the local Maxima defines the intensity of the clusters. For each nanocluster a 2D Gaussian equation was fitted in a 5 by 5 pixels patch centred on the Maxima coordinates, in x and y. FWHM in x and y were averaged to get the mean FWHM reported in the text. The data was then processed using Matlab (using the script provided) and Microsoft Excel. For the colocalization of Arc nanoclusters with another marker (Arc mRNA, AMPA receptors or PSD95), the STED images in the channel of the marker were analysed using the script provided. A Laplatian filtering (smoothing: 3) and a Gaussian Blurring (sigma: 1) were first applied to the images, each image was then auto-thresholded and binarized. In binary images the Watershed and morphological (radius: 1) filters were applied and just particles larger than 0.01 um were detected and enlarged by 0.05 um. The areas identified are the areas in which local Arc maxima are found in Arc channel images. Arc nanoclusters within defined areas (colocalization with Arc) and outside (extrasynaptic Arc nanoclusters) are separately analysed using the same script as reported above in order to identify the size of Arc nanoclusters which is, or which are not co-localizing with the other marker. For the identification of EZ, PSD95 areas were radially enlarged of 200 nm, each PSD95 area was then subtracted from its enlarged counterpart, Arc local maxima found in the resulting area were attributed to the EZ. The percentage of arc nanoclusters in colocalization with Arc mRNA is calculated dividing the numbers of clusters in colocalization over the total number of nanoclusters detected in STED imaging.

The line profiles were traced using smart neighborhood of three pixels with the software Imspector (v0.1) and fitted with a GaussAmp function using OriginLab software.

### DNA-PAINT Image Analysis

Raw DNA-PAINT data was reconstructed to super-resolution images with the Picasso software package (version 0.6.0)^70^, latest version available at https://github.com/jungmannlab/picasso). Drift correction was performed with a redundant cross-correlation following gold particles as fiducials for cellular experiments. Alignment of sequential imaging rounds was performed using gold particles.

Arc, Clathrin and PSD95 clusters of localizations were obtained using the DBSCAN algorithm^71^. DBSCAN parameters, minimum localizations, and clustering radius were selected based on the imaging parameters of the individual super-resolution channel of the protein. Background regions were taken as a base for determining the cluster minimum localization number and both DBSCAN parameters were further adjusted based on visual validation to determine a cutoff value distinguishing background from specific protein clusters. Cluster volumes are calculated by finding a 3D convex hull and extracting its volume. Distances between clusters of different protein types were calculated subsequently.

Quantification of Arc accumulation at the synaptic level was obtained from DNA-PAINT images measuring the local density of Arc single molecules (counted with ImageJ local maxima finder) in colocalization with PSD95 areas in comparison to extra synaptic Arc densities. Just the synapses where Arc single molecules were 30% more abundant than in extrasynaptic areas were considered as synapses with Arc accumulations. Arc molecules to PSD95 nanomodules alignment was quantified upon tracing 120 nm averaged line profiles along PSD from axially sliced and 65 nm y-projected images of top-viewed synapses.

## STARSS Data Analysis and Statistics

A mono-exponential fitting function was used to model the data,

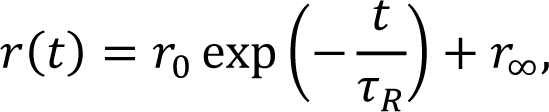

where 𝑟 is the fluorescence anisotropy computed from the signals recorded in the parallel and perpendicular channels, 𝑡 is time, 𝜏_𝑅_ is the rotational diffusion time, 𝑟_0_ is the anisotropy that decays with time constant 𝜏_𝑅_ in the observed time window, and 𝑟_∞_ is the anisotropy fraction that does not decay in the observed time window.

Signals from 56 field of views of 20×20 micrometer squared, recorded from 28 neurons derived from 3 independent cultures in four sessions in different days, were manually segmented in spines and dendrites. The signals were spatially integrated and summed together to obtain time decays representative of the full dataset. Errors are computed propagating the photon shot noise of the detectors in the anisotropy formula^23^. All the FOV with total counts less than 3×10^4 were discarded. For the discarded FOV the confidence interval computed by propagating the photon shot noise was already high, and the background subtraction was challenging and inducing unwanted biases. This estimation of the structure size of the fraction of anchored protein is obtained by converting the rotational diffusion time to the hydrodynamic diameter using the Stokes-Einstein equation^23^, assuming room temperature and viscosity of water. The fitting parameters are reported in the following table.

| Segmentation | $\tau_R$ (us) | $d_R$ (nm) | $r_0$ | $r_\infty$ |
| --- | --- | --- | --- | --- |
| Dendrites | 29.6 | 61 | 0.011 | 0.0069 |
| Spines | 88.6 | 88 | 0.010 | 0.0070 |

For statistical significance of anisotropy decays in dendritic spines and shaft we applied the modified Chi-squared method reported by Hristova et al., 2023^72^ to measure differences between arbitrary curves for the scenario of a single measurement at each point with known standard deviation.

## Statistical Analysis

The statistical tests applied to compare pooled data from multiple replicas were the two-sample two-sided Student’s t-test for normally distributed data and the non-parametric Kolmogorov– Smirnov test for non-normally distributed data.

