## Supplementary Information for "Structural diversity of Arc oligomers within the excitatory synapse"

### List of content

#### Supplementary notes

1. Cloning design and constructs
2. Immunostaining in HeLa cells

#### List of Figures

1. Arc is organized in nanoclusters in primary neuronal cultures.
2. Arc nanoscale organization at the excitatory synapse.
3. Arc oligomers form liquid condensates in cells.
4. Palmitoylation and PIP lipids modulate but they are not required to mediate Arc-membrane interaction.
5. Arc induces membrane inward bending and high-order oligomers affect AMPA receptors surface levels.

#### List of Tables

1. Arc mRNA FISH probes
2. DNA-PAINT imaging parameters

**Supplementary notes**

**Supplementary Note 1: Cloning design and constructs**

- pAAV-EF1a_Arc-FL-C-SNAP-IRES-WGA-Cre was generated by first exchanging mCherry from p83_pAAV-EF1a-mCherry-IRES-WGA-Cre (kindly provided by Michael Ratz from Department of Cell and Molecular Biology, Karolinska Institutet) with SNAP amplified from the plasmid template Sec61β-SNAP (kindly provided by Francesca Bottanelli, Freie Universität Berlin) using the primer pair

5′- atgactggatccATGGACAAAGACTGCGAAATGAAGC

5'- agtcatgaattcCTAACCCAGCCCAGGCTTGCCC

Both backbone and insert were digested using BamHI-HF and EcoRI-HF (NEB) separately, followed by T4 ligation (NEB) and transformation of NEB Stable cells (NEB). Then, rat Arc was amplified from pcDNA3.1(+)-C-eGFP-ARC using the primer pair.

5'- atgactggtaccggattggccaccATGGAGCTGGACCATATGACG

5'- agtcatggatccgctaccgctgccTTCAGGCTGGGTCCTGTCACT

Both backbone and insert were digested using BamHI-HF and KpnI-HF (NEB) separately, followed by T4 ligation (NEB) and transformation of NEB Stable cells (NEB).

- pAAV-EF1a_Arc-IM-fusion-SNAP-IRES-WGA-Cre was generated digesting the backbone pAAV-EF1a_Arc-FL-C-SNAP-IRES-WGA-Cre using NcoI (NEB) and amplifying three sequences from it using the following three pairs of primers:

ARC_CTD_rev_ g: 5'- tatcgataagcttgatatcgttaTTCAGGCTGGGTCCTGTC

ARC_NTD_fwd_g: 5'- tcaggtgtcgtgaggtaccgggtttagtgaaccgtGCCACCATGGAGCTGGACC

ARC_NTD_rev_SNAP: 5'- ttcatttcgcagtctttgtcgccgccgatggagccGGACTCCAGGCGGTCGGC

ARC_SNAP_fwd: 5'- gggccgaccgcctggagtccggctccatcggcggcGACAAAGACTGCGAAATGAAGCGCAC

ARC_SNAP_rev: 5'- actgggtacttgccgcccatgatggagccgccACCCAGCCCAGGCTTGCC

ARC_CTD_fwd_SNAP: 5'- tgggcaagcctgggctgggtggcggctccatcATGGGCGGCAAGTACCCA

Backbone and PCR products were assembled with Gibson Assembly (Master Mix, NEB).

- pAAV-EF1a_Arc-linker-CTD-SNAP-IRES-WGA-Cre was generated from pAAV-EF1a_Arc-FL-C-SNAP-IRES-WGA-Cre digesting the backbone with NcoI (NEB) followed by T4 ligation (NEB) and transformation of NEB Stable cells (NEB).

- pAAV-EF1a_Arc-linker-CTD-SNAP-IRES-WGA-Cre was generated from pAAV-EF1a_Arc-FL-C-SNAP-IRES-WGA-Cre digesting the backbone with NcoI (NEB) and BamHI-HF (NEB) and amplifying from it a sequence using the following primers pair:

5'- tgaggtaccggattggccacCCATGGAGCTGGACCATATGAC

5'- tcatttcgcagtctttgtcgggatccgctaccgctgccCCATGGACTCCAGGCGGT

The backbone and PCR product were assembled with Gibson Assembly (Master Mix, NEB) and and transformed with NEB Stable cells (NEB).

- pAAV-EF1a_ArcC94-98S-C-SNAP-IRES-WGA-Cre was generated from pAAV-EF1a_Arc-FL-C-SNAP-IRES-WGA-Cre with Site directed mutagenesis (Q5® Site-Directed Mutagenesis Kit, NEB) using the following primers pair:

5'- CCAGTCTCAGCCGCAGCCAGGAGACCATCGCCAACC

5'- CCTTGATGGACTTCTTCCAGCG

- pAAV-EF1a_ArcC94-98A-C-SNAP-IRES-WGA-Cre was generated from pAAV-EF1a_Arc-FL-C-SNAP-IRES-WGA-Cre with Site directed mutagenesis (Q5® Site-Directed Mutagenesis Kit, NEB) using the following primers pair:

5'- cGgcGgcGGcGGcCTTCTACCGTCTGGAGAGGTGGG

5'- cCgcCgCGgcCgcCTCACGCTTGACCCAGCGCTC

- pAAV-EF1a_Arc-FL-C-rsEGFP2-IRES-WGA-Cre was generated by first exchanging mCherry from p83_pAAV-EF1a-mCherry-IRES-WGA-Cre (kindly provided by Michael Ratz from Department of Cell and Molecular Biology, Karolinska Institutet) with rsEGFP2 amplified from the plasmid template rsEGFP2-Omp25[73] using the primer pair

5'- atgactggatccATGGTGAGCAAGGGCGAGGAG

5'- agtcatgaattcTCACTTGTACAGCTCGTCCATGCCG

Both backbone and insert were digested using BamHI-HF and EcoRI-HF (NEB) separately, followed by T4 ligation (NEB) and transformation of NEB Stable cells (NEB). Then, rat Arc was amplified from pcDNA3.1(+)-C-eGFP-ARC using the primer pair.

5'- atgactggtaccggattggccaccATGGAGCTGGACCATATGACG

5'- agtcatggatccgctaccgctgccTTCAGGCTGGGTCCTGTCACT

Both backbone and insert were digested using BamHI-HF and KpnI-HF (NEB) separately, followed by T4 ligation (NEB) and transformation of NEB Stable cells (NEB).

- pAAV-EF1a_Arc-IM-fusion-rsEGFP2-IRES-WGA-Cre was generated digesting the backbone pAAV-EF1a_Arc-FL-C-rsEGFP2-IRES-WGA-Cre using NcoI (NEB) and amplifying three sequences from it using the following three pairs of primers:

ARC_CTD_rev_ g: 5'- tatcgataagcttgatatcgttaTTCAGGCTGGGTCCTGTC

ARC_NTD_fwd_g: 5'- tcaggtgtcgtgaggtaccgggtttagtgaaccgtGCCACCATGGAGCTGGACC

ARC_NTD_rev_rs2: 5'- agctcctcgcccttgctcac gccgccgatggagccgGACTCCAGGCGGTCGGCC

ARC_rsEGFP2_fwd: 5'- gggccgaccgcctggagtccggctccatcggcggcGTGAGCAAGGGCGAGGAG

ARC_rsEGFP2_rev: 5'- actgggtacttgccgcccatgatggagccgccCTTGTACAGCTCGTCCATGC

ARC_CTD_fwd_rs2: 5'- gcatggacgagctgtacaagggcggctccatcATGGGCGGCAAGTACCCA

The backbone and PCR products were assembled with Gibson Assembly (Master Mix, NEB).

- pSFV-SCA_5'-UTR_Arc-IM-fusion-SNAP_3'UTR was generated from Semliki-Forest Virus pSCA3-LifeAct-DronpaM159T (kindly gifted by Dr. Stefan W. Hell, MPI-BCP Göttingen, Germany) which was digested with XmaI-HF and NotI-HF (NEB). Arc-IM-fusion-SNAP sequence was amplified from pAAV-EF1a_Arc-IM-fusion-SNAP-IRES-WGA-Cre using the following primers pair:

5'- cggcgagcagATGGAGCTGGACCATATGAC

5'- gctggcccctTTATTCAGGCTGGGTCCTG

rat Arc-UTR and 3'-rat Arc-UTR sequences were amplified from costumed gene fragment cDNA purchased from IDT (Integrated DNA Technologies) using the two following couples of primers:

5'- cacagaattctgattggatcAGTGCTCTGGCGAGTAGTCC

5'- ccagctccatCTGCTCGCCGGGGTTACG

5’- gcctgaataaAGGGGCCAGCCCAGGGTC

5’-
aattcaattaattaccctgcTTTAAATTTCATAGTTTTATTAACAAAATCATATATATGTATATAT
ATATATATGTCTGTCTTTGAGGTAAGATGGTGTGGGCCAGATGG

The backbone and PCR products were assembled with Gibson Assembly (Master Mix, NEB).

- pSFV-SCA_Arc-FL-C-AlfaTag was generated from Semliki-Forest Virus pSCA3-LifeAct-DronpaM159T (kindly gifted by Dr. Stefan W. Hell, MPI-BCP Göttingen, Germany) which was digested with XmaI-HF and NotI-HF (NEB). Arc sequence was amplified from pAAV-EF1a_Arc-FL-C-SNAP-IRES-WGA-Cre using the following primers pair:

5’- cacagaattctgattggatcGCCACCATGGAGCTGGAC

5’- GGggatccgctaccgctgccttcaggctgggtcctgtcac

AlfaTag sequence is made by the annealing of two primers. Backbone and products were assembled with Gibson Assembly (Master Mix, NEB).

**Immunostaining in HeLa cells**

For the immunostaining of untagged Arc-FL or Arc-FL-C-EGFP and G3BP1 protein (marker for stress granules) in HeLa cells, cells were washed once with 1X PBS, pre-warmed at 37°C and fixed in pre-warm 4% PFA at room temperature for 20 minutes. After being washed twice with PBS, cells were permeabilize by incubation with 0.5% TRITON™ X100 in PBS for 5 minutes or 0.1% TRITON™ X100 in PBS for 10 minutes. Blocking was performed by incubation with 5% bovine serum albumin in PBS (5% BSA/PBS) for 30 minutes at RT. Anti-Arc Antibody (C-7, sc-17839, Santa Cruz, 200 µg/ml, used 1:50) or Anti-G3BP1 (mouse, kindly gifted by Marc Panas, Karolinska Institutet) were incubated in 5% BSA/PBS for 1 hour at RT. After three 5 min washes in PBS, the sample was incubated with donkey anti-mouse Alexa594 secondary antibody (ThermoFisher A-21203, 2 mg/ml, 1:200) and with Anti-GFP nanobody (FluoTag®-X4 anti-GFP conjugated with Abberrior635P, N0304, 1:500) in 5% BSA/PBS for 1 hour at RT. The samples were washed in PBS for 15 min and the coverslips were mounted I Mowiol.

**Supplementary Figures**

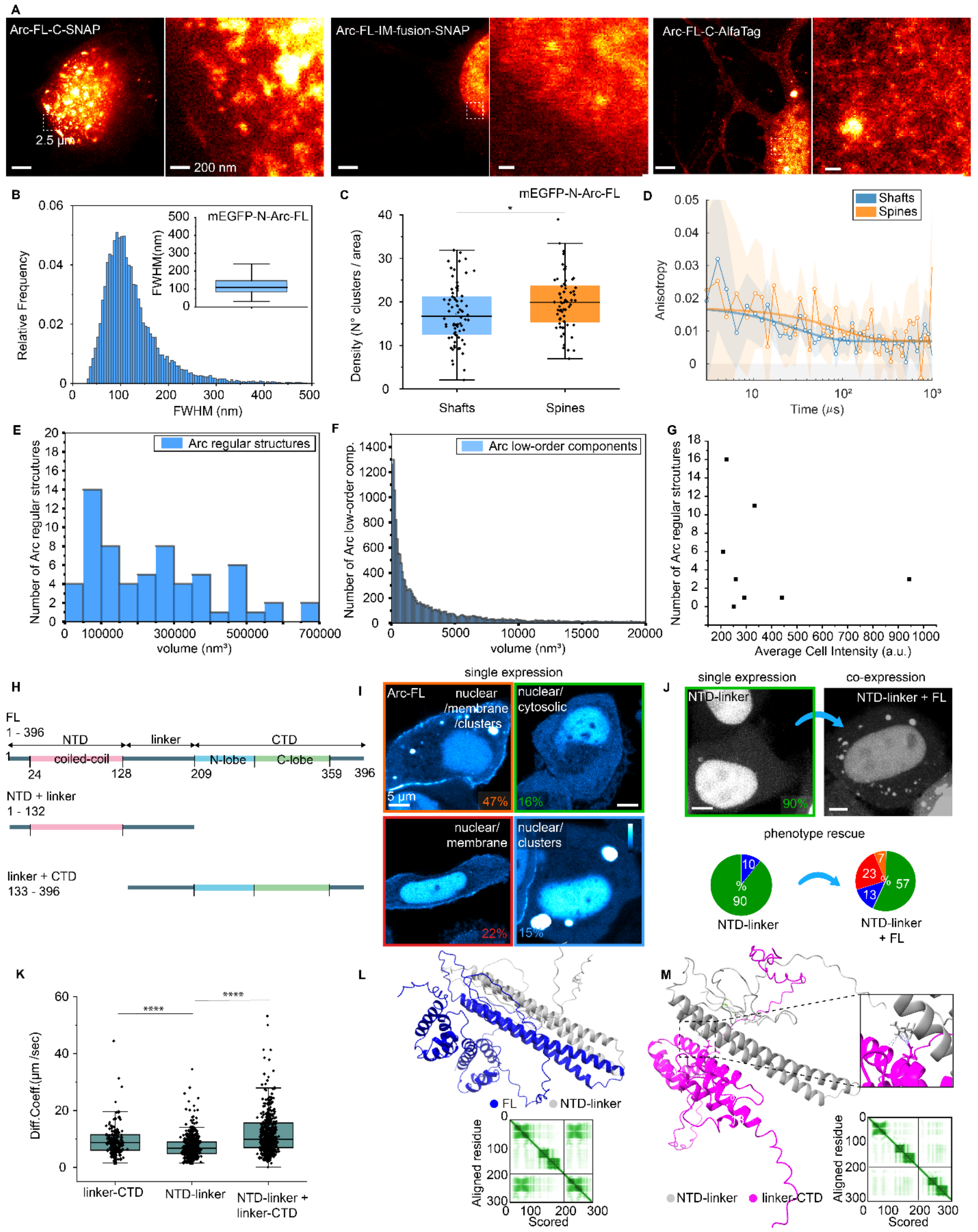


**Supplementary Fig. 1. Arc is organized in nanoclusters in primary neuronal cultures.**

(A) Immature cortical neurons (DIV7-10) showing Arc localization in the nucleus for the different Arc labeling strategies: Arc-FL-C-SNAP (left panels), Arc-FL-IM-fusion-SNAP (central panels), Arc-FL-C-AlfaTag (right panels).

(B) The histogram reports mEGFP-N-Arc-FL nanocluster FWHM distribution obtained from 23 neurons (DIV17-21) derived from 5 samples in 2 independent cultures.

(C) The box plot represents the significantly higher density of mEGFP-N-Arc-FL nanoclusters in spines versus dendritic shaft. Each data plot is the average density value per spines and shaft per image. Two-sample two-sided Student's t test p-value: 0.01708, the box plots show the 25–75% interquartile range, with the middle line representing the mean, and the whiskers derived from 1.5 * interquartile range t test p-value).

(D) Anisotropy decays measured with STARSS and mono-exponentially fitted for Arc-FL-IM-fusion-rsEGFP2 in spines (orange) and in shafts (blue).

(E) The histogram reports the volume of Arc regular structures (high-order oligomers) distribution as measured in 3D DNA-PAINT from 21 cortical neurons (DIV21-22) derived from 2 independent cultures.

(F) The histogram reports the volume of Arc low-order oligomers distribution as measured in 3D DNA-PAINT from 21 cortical neurons (DIV21-22) derived from 2 independent cultures.

(G) The scatter plot reports the lack of correlation between the number of detected regular structures and the overall mEGFP-N-Arc-FL cell expression level.

(H) Schematic representation of three Arc variants used: Arc-FL (1 - 396 aa, NTD-linker-CTD), Arc NTD-linker (1 - 132 aa), Arc linker-CTD (133 – 396 aa).

(I) Representative images of HeLa cells exogenously expressing full-length Arc. Identified phenotypes: nuclear, plasma membrane, and cytosolic localization with bright clusters (orange, 47%, N=67 cells), homogeneous cytosolic and nuclear localization (green, 16%, N=22 cells), homogenous cytosolic, nuclear and plasma membrane localization (red, 22%, N=32 cells), nuclear and cytosolic with bright clusters (blue, 15%, N=21 cells).

(J) (upper left panel) HeLa cells exogenously expressing Arc NTD-linker. (upper right panel) Arc NTD-linker channel of HeLa cells exogenously co-expressing Arc-FL and NTD-linker. Lower right pie chart: NTD-linker identified phenotypes, cytosolic homogeneous and nuclear (90%) nuclear and homogenous cytosolic with bright clusters (10%). Lower right pie chart: phenotype rescue observed for Arc NTD-linker when co-expressed with Arc-FL, cytosolic homogenous and nuclear (57%, N=17 cells), nuclear and homogenous cytosolic with bright clusters (13%, N=4), plasma membrane localization together with cytosolic clusters, (7%, N=3) and plasma membrane localization (23%, N=11).

(K) The plot shows the significant difference among diffusion coefficients ($D_t$) for Arc linker-CTD, NTD-linker, both in single expression, and NTD-linker in co-expression with rat Arc linker-CTD in HeLa cells. (CTD-linker median $D_t$: 8,71 $\mu m^2/s$, linker-NTD median $D_t$: 6,73 $\mu m^2/s$, NTD-linker + linker-CTD median $D_t$: 9,89 $\mu m^2/s$, box plots show the 25–75% interquartile range, with the middle line representing the mean, and the whiskers derived from 1.5 * interquartile range. Two-sample two-sided Kolmogorov–Smirnov test p-values respectively: $3.8 \times 10^{-5}$ and $1.8 \times 10^{-21}$).

(L) AlphaFold v2.1.0 Multimer interaction prediction Arc-FL (blue) and Arc NTD-linker (grey) with the expected position error plot (pLDDT model confidence).

(M) AlphaFold v2.1.0 Multimer interaction prediction for Arc NTD-linker (grey) and linker-CTD (magenta) with the expected position error plot (pLDDT model confidence). Inset: major sites of protein-protein interactions in between the first α-helix of the NTD and CTD N-lobe.

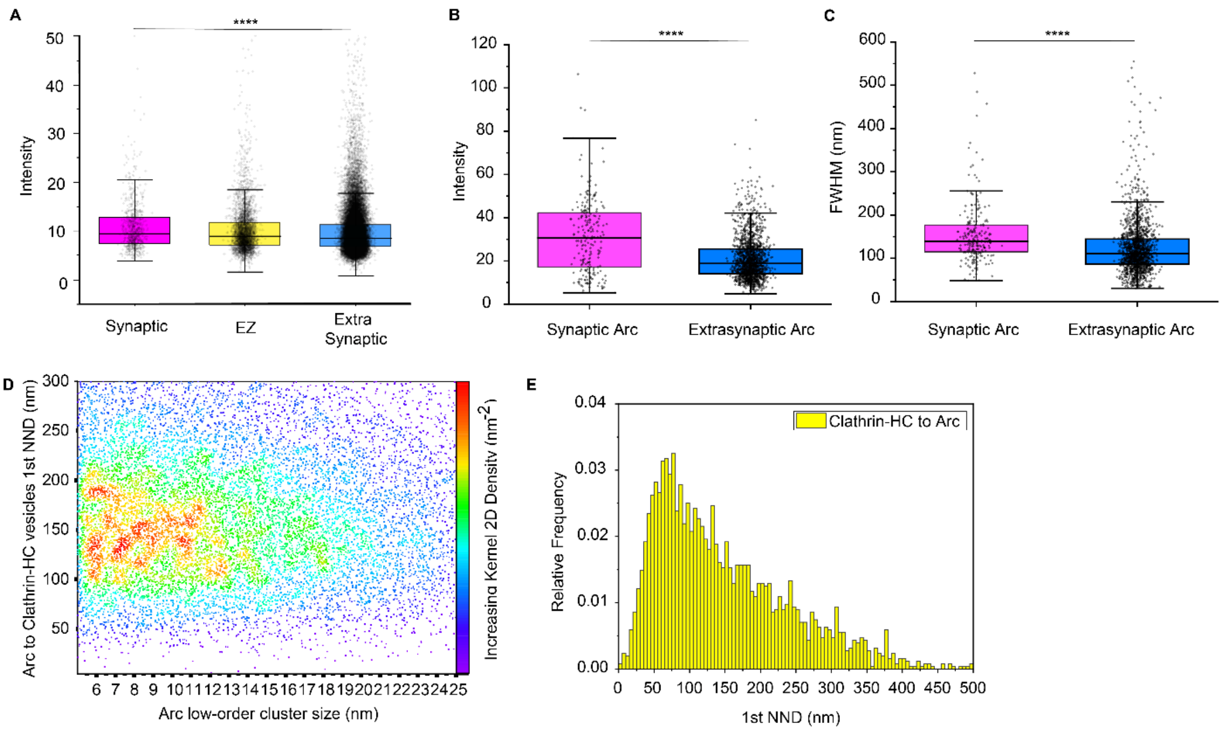

**Supplementary Fig. 2. Arc nanoscale organization at the synaptic compartment.**

(A) The box plot of immunolabelled endogenous Arc shows significant difference in brightness between Synaptic Arc nanoclusters (co-localizing with PSD95), Arc within the EZ (within 200 nm from PSD95 perimeter) and extrasynaptic Arc from 26 neurons derived from 3 independent cultures. The box plots show the 25–75% interquartile range, with the middle line representing the mean, and the whiskers derived from 1.5 * interquartile range. Two-sample two-sided Kolmogorov–Smirnov test p-values respectively: p = 5.6487e-10, p = 2.1644e-04, p = 1.2136e-06).

(B) The box plot shows the significant difference in brightness between mEGFP-N-Arc-FL nanoclusters co-localizing with PSD95 (Synaptic Arc) and extrasynaptic Arc derived from the neuron shown in Fig. 2B. The box plots show the 25–75% interquartile range, with the middle line representing the mean, and the whiskers derived from 1.5 * interquartile range. Two-sample two-sided Kolmogorov–Smirnov test p-value: 1.1010e-25.

(C) The box plot shows the significant difference in size between mEGFP-Arc nanoclusters co-localizing with PSD95 (Synaptic Arc) and extrasynaptic Arc derived from the neuron shown in Fig. 2B. The box plots show the 25–75% interquartile range, with the middle line representing the mean, and the whiskers derived from 1.5 * interquartile range. Two-sample two-sided Kolmogorov–Smirnov test p-value: 1.0575e-15.

(D) The scatter plot shows the relationship between Arc to clathrin-coated vesicles 1$^{st}$ NNDs to Arc, the density color map for each datapoint reveals several higher density areas corresponding to different Arc low-order cluster sizes located within 100 nm to 200 nm from clathrin-coated vesicles.

(E) The histogram reports the distributions of 1$^{st}$ NNDs from clathrin-coated vesicles to Arc which peaks at 75 nm.

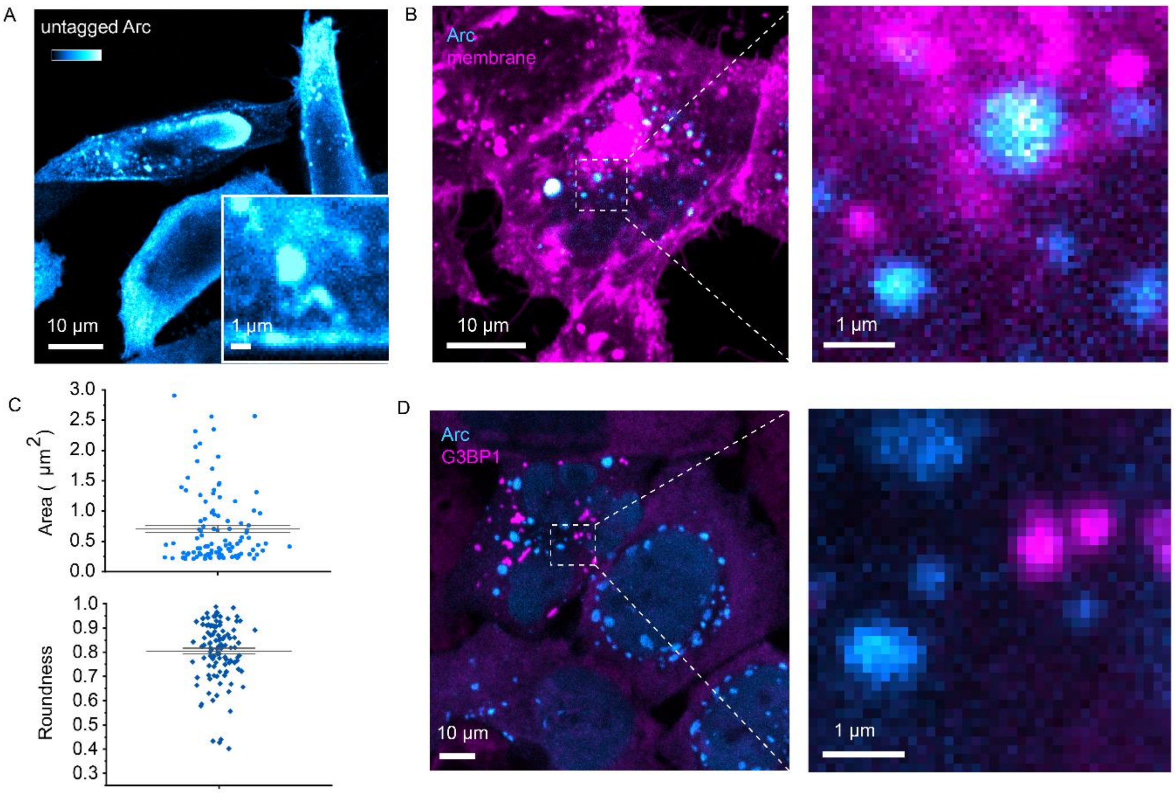


**Supplementary Fig. 3. Arc oligomers form liquid condensates in cells.**

(A) Representative image of untagged Arc-FL expressed in HeLa cells and immunostained. Phenotype with cytosolic bright microclusters is observed.

(B) Representative image of HeLa cells exogenously expressing Arc-FL (C-EGFP tagged) where membranes were labelled with a lipophilic dye to show that there is no co-localization between Arc micrometric condensates and membrane-bound intracellular organelles such as lysosomes (zoom-in).

(C) (Upper panel) scattered interval plot where distribution of Arc clusters area is presented (average area: 0.705 $um^2$). (Lower panel). Scattered interval plot where distribution of Arc clusters roundness is presented (average roundness 0.8). For both plots it is shown the 25–75% interquartile range, with the middle line representing the mean, and the whiskers are derived from the standard error.

(D) Representative image of HeLa cells exogenously expressing Arc-FL (C-EGFP tagged) and immunostained for G3BP1 protein to show that there is no co-localization between Arc micrometric condensates and stress granules.

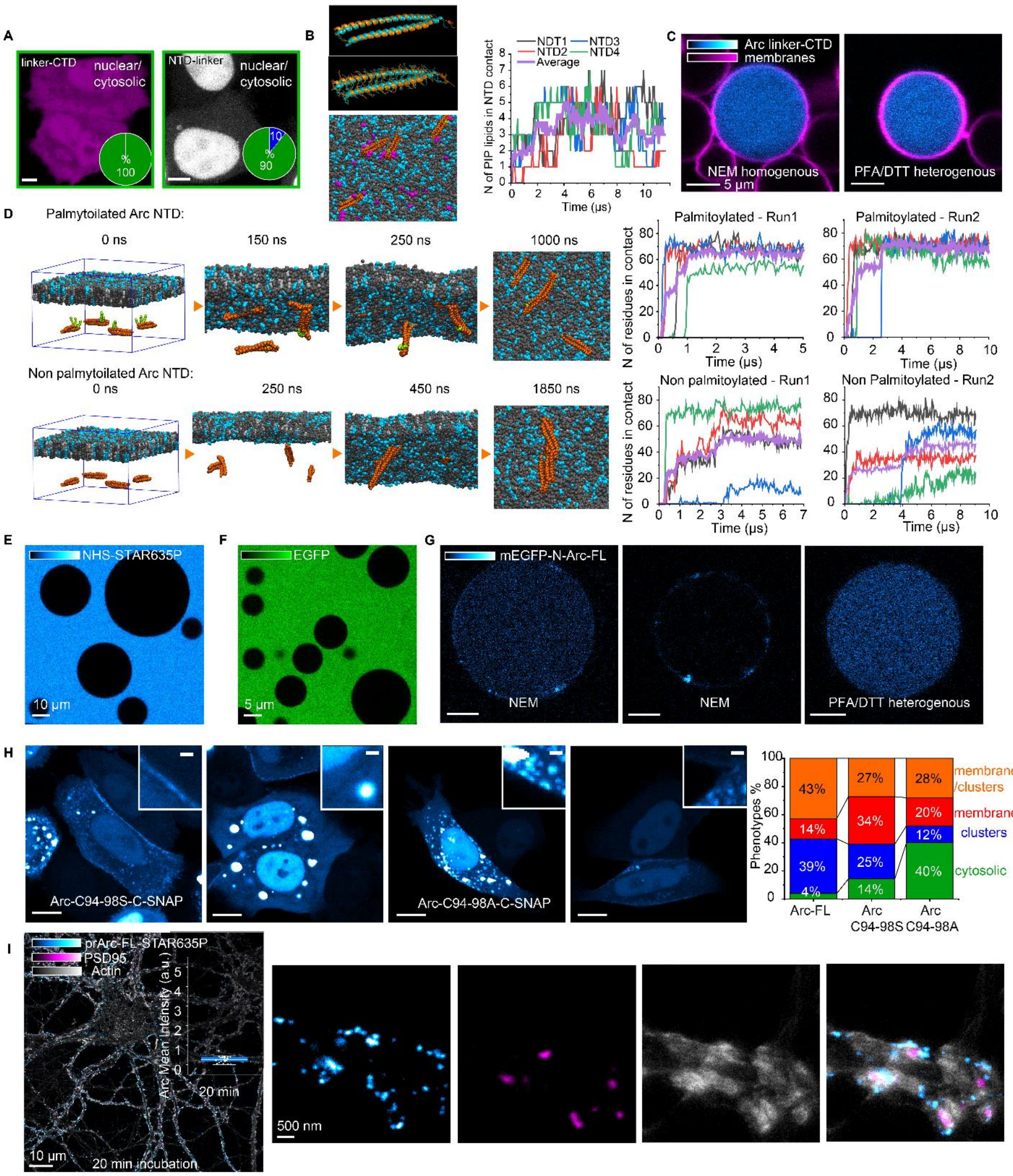


**Supplementary Fig. 4. Palmitoylation and PIP lipids modulate but they are not required to mediate Arc-membrane interaction.**

(A) (Left) Representative image of HeLa cells exogenously expressing Arc linker-CTD. The single expression phenotypes are reported in the inset pie chart: 100% cytosolic homogenous. (Right) Representative Arc NTD-linker channel image of HeLa cells exogenously expressing Arc NTD-linker. The single expression phenotypes are reported in the inset pie chart: 90% cytosolic homogenous and 10% cytosolic microclusters. Scale bar: 5 µm.

(B) Comparison between the NTD of rat Arc from AlphaFold (orange) and TrRosetta (teal). The top and bottom images show the same structures, but the bottom image omits the individual residues from the representation. The two helices of both structures are well aligned with RMSD of 3.36 Å (CE alignment) or 2.72 Å (SALIGN), as computed in PyMod plugin in PyMol.

(C) (left panel) Increased density of PIP lipids (magenta) in proximity of NTDs (orange) at time = 11.8 ⌠s from the beginning of the simulation. Note that PIP lipids do not bind permanently to the NTDs but are able to detach and diffuse further along the membrane. (right panel) The number of PIP lipids bond vs. time for each of the 4 NTDs (thin colored lines) and the average over all NTDs (thick violet line).

(D) Two examples of NEM-produced GPMVs derived from Arc linker-CTD expressing HeLa cells, showing homogeneous Arc layout and no Arc micrometric clusters.

(E) Two coarse-grained molecular dynamics systems: (upper panel) palmitoylated NTDs and (lower panel) non-palmitoylated NTDs. In the palmitoylated NTDs simulation, two NTDs interact in solution with each other and one with the membrane at 150 ns. At 250 ns two of the NTDs are bound to the membrane. At 1000 ns, all four NTDs are bound to the membrane. In the non-palmitoylated NTDs simulation two of the NTDs interact in the solution at 250 ns. At 450 ns, the third NTD interacts with the first two, whereas the fourth binds to the membrane. The three interacting NTDs attach to the membrane at 1850 ns. The graphs on the right quantify the number of NTD residues in contact with the lipids for each of the NTDs and their average (thick violet line). The result was confirmed by two sets of simulations.

(F) GUVs incubated with NHS-STAR635P showing absence of the interaction between the dye and the GUVs lipid bilayers.

(G) GUVs incubated with EGFP showing absence of the interaction between the protein and the GUVs lipid bilayers.

(H) (left and central panel) Examples of NEM-produced GPMVs derived from mEGFP-N-Arc-FL expressing primary cortical neurons (DIV21) showing comparable results to HeLa cells derived GPMVs: Arc micrometric clusters in proximity and co-localizing with the GPMVs membranes. (right panel) Example of PFA/DTT-produced GPMVs derived from mEGFP-N-Arc-FL expressing primary cortical neurons (DIV21) showing comparable results to HeLa cells derived GPMVs: no homogeneous Arc layout and no resolved Arc micrometric clusters.

(I) HeLa cells exogenously express Arc-C94-96S and C94-98A-SNAP. The plot represents the different phenotype percentages observed for Arc-FL compared to Arc-C94-96A and Arc-C94-98S mutants: compared to Arc-FL a phenotype shift is observed for the cytosolic homogeneous (green) component, from 4% to 14% (N=11) for ArcC94-98S, and to 40% (N=10) for ArcC94-98A.

(J) Representative primary cortical neurons (DIV21) incubated for 20 min with rat prArc-FL-STAR635P (4 ug) and immunostained against PSD95 and labelled for actin. Arc is shown in interaction with extracellular leaflet of neuronal plasma membrane accumulating at the level of dendritic spines. In the inset, the box plot shows the variability of Arc mean intensities for different neurons.

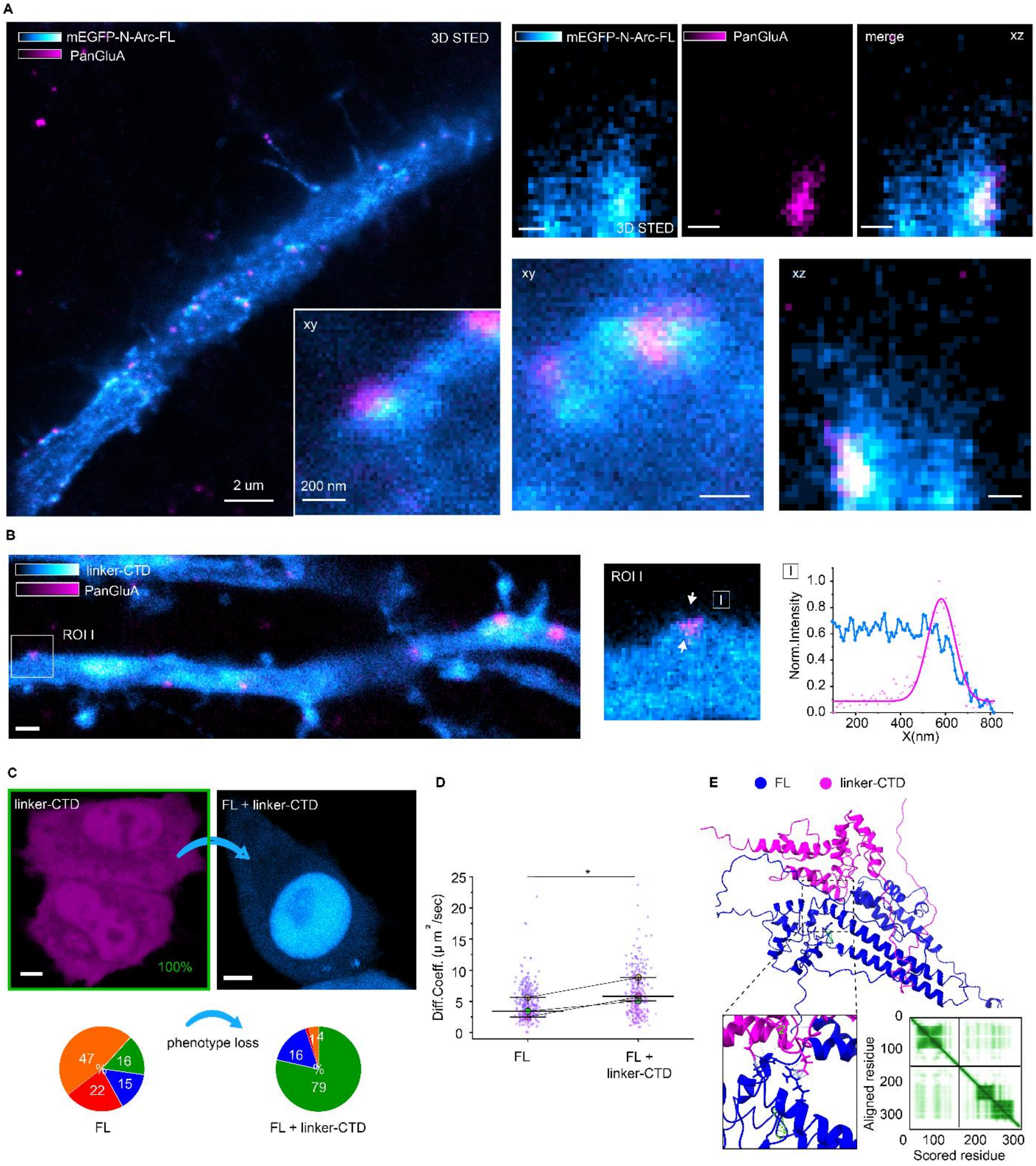


**Supplementary Fig. 5. Arc induces membrane inward bending and high-order oligomers affect AMPA receptors surface levels.**

(A) Mature hippocampal neurite (DIV21) expressing mEGFP-N-Arc-FL (cyan) and PanGluA (magenta), live stained prior to fixation and imaged in 3D STED microscopy. Arc nanoclusters colocalization persists in 3D STED showing the independence of Arc colocalization from the geometry of the compartment.

(B) Mature cortical neurite (DIV20-21) expressing Arc linker-CTD-SNAP (cyan) and PanGluA (magenta), live stained prior to fixation and imaged in STED microscopy. In ROI I, no Arc nanoclusters are seen in co-localization with GluA, line profile traced along the white arrows.

(C) (Upper left panel) HeLa cells exogenously express Arc linker-CTD with cytosolic homogeneous phenotype (100%). (Upper right panel) Arc-FL channel of HeLa cells in co-expression with Arc linker-CTD. Observed phenotype loss for Arc-FL from single expression (left pie chart) reported in the lower right pie chart.: nuclear and cytosolic homogenous localization (79%, N=62), nuclear cytosolic homogenous with bright clusters (16%, N=13), plasma membrane localization together with cytosolic clusters (4%, N=3), membrane localization (1%, N=1).

(D) The plot shows the significant difference between $D_t$ for Arc-FL when expressed alone or in co-expression with Arc linker-CTD. FL median $D_t$: 4.9 $\mu m^2/s$, FL+ linker-CTD median $D_t$: 7.3 $\mu m^2/s$). The plots' the middle line represents the mean, and the whiskers are derived from 1.5 * interquartile range. Two-sample two-sided Kolmogorov–Smirnov test p-value: 0.0346, from 3 independent experiments whose mean values are color-coded.

(E) AlphaFold v2.1.0 Multimer interaction prediction for Arc-FL (blue) and linker-CTD (magenta) with the expected position error plot (pLDDT model confidence). Inset: major sites of protein-protein interactions resulting from disruption of Arc-FL structure. Most of the contacts occur between the linker-CTD, at the level of the hinge region connecting N-lobe and C-lobe ($_{146}$TLSREA$_{151}$), and FL C-lobe. These interactions induced conformational changes in the 3D structure of Arc-FL in between 4 residues ($_{216}$DPREP$_{220}$) at the N-lobe and Glu118 within the motif ($_{113}$MHVWREV$_{119}$), which was shown to be required for oligomerization above the dimeric state [14].

| | | |
|---|---|---|
| | ACGTAGGATCGTTGCTCGTC ACGTAGGATCGTTGCTCGTC | |
| Consensus | ACGTAGGATCGTTGCTCGTC------------------------------ACGTAGGATCGTTGCTCGTC |  |
| oligo_6 [rn6]chrArc:217-247 | ACGTAGGATCGTTGCTCGTCGCCGCCGGTCGTCATATGGTCCAGCTCCATACGTAGGATCGTTGCTCGTC | 70 |
| oligo_7 [rn6]chrArc:250-280 | ACGTAGGATCGTTGCTCGTCCGGCCCACCCCGCGGGGCAGGGTAGGCGTGACGTAGGATCGTTGCTCGTC | 70 |
| oligo_8 [rn6]chrArc:288-318 | ACGTAGGATCGTTGCTCGTCCACTTACCAATCTGCAGGATCACATTGGGTACGTAGGATCGTTGCTCGTC | 70 |
| oligo_9 [rn6]chrArc:321-351 | ACGTAGGATCGTTGCTCGTCGTCCTCCGTACGTGCTCCAGCATCTCAGCTACGTAGGATCGTTGCTCGTC | 70 |
| oligo_10 [rn6]chrArc:352-382 | ACGTAGGATCGTTGCTCGTCCTTGGACACTTCGGTCAACAGATGCCGGTGACGTAGGATCGTTGCTCGTC | 70 |
| oligo_11 [rn6]chrArc:395-425 | ACGTAGGATCGTTGCTCGTCTGCCCACCGACCTGTGCAACCCTTTCAGCTACGTAGGATCGTTGCTCGTC | 70 |
| oligo_12 [rn6]chrArc:428-458 | ACGTAGGATCGTTGCTCGTCTGGGCACATAGCCGTCCAAGTTGTTCTCCAACGTAGGATCGTTGCTCGTC | 70 |
| oligo_13 [rn6]chrArc:461-491 | ACGTAGGATCGTTGCTCGTCTGATGGACTTCTTCCAGCGCTGTGAGTCGCACGTAGGATCGTTGCTCGTC | 70 |
| oligo_14 [rn6]chrArc:498-528 | ACGTAGGATCGTTGCTCGTCTTGGCGATGGTCTCCTGGCAGCGGCAGAGAACGTAGGATCGTTGCTCGTC | 70 |
| oligo_15 [rn6]chrArc:530-560 | ACGTAGGATCGTTGCTCGTCCGTGCATCTCACGCTTGACCCAGCGCTCCAACGTAGGATCGTTGCTCGTC | 70 |
| oligo_16 [rn6]chrArc:561-591 | ACGTAGGATCGTTGCTCGTCCTCTCCAGACGGTAGAAGACCTCCCTCCACACGTAGGATCGTTGCTCGTC | 70 |
| oligo_17 [rn6]chrArc:592-622 | ACGTAGGATCGTTGCTCGTCGCCGCCCATGGACTCCAGGCGGTCGGCCCAACGTAGGATCGTTGCTCGTC | 70 |
| oligo_18 [rn6]chrArc:630-660 | ACGTAGGATCGTTGCTCGTCACAGTGTGGCGGGCCGGCTCGCTGCCCACTACGTAGGATCGTTGCTCGTC | 70 |
| oligo_19 [rn6]chrArc:665-695 | ACGTAGGATCGTTGCTCGTCGGCAGTAGGGCTCTGGACCCCCCACACCTAACGTAGGATCGTTGCTCGTC | 70 |
| oligo_20 [rn6]chrArc:697-727 | ACGTAGGATCGTTGCTCGTCGCTAACAGTGTAGTCGTAGCCATCAGCTTCACGTAGGATCGTTGCTCGTC | 70 |
| oligo_21 [rn6]chrArc:752-782 | ACGTAGGATCGTTGCTCGTCCTGACTCCTGCTCAGGCAGCTCTCCTGCGGACGTAGGATCGTTGCTCGTC | 70 |
| oligo_22 [rn6]chrArc:783-813 | ACGTAGGATCGTTGCTCGTCGGCACCCAAGACTGGTATTGCTGAGCCCCAACGTAGGATCGTTGCTCGTC | 70 |
| oligo_23 [rn6]chrArc:814-844 | ACGTAGGATCGTTGCTCGTCCAGACCTGGGCTTGGTTGCCCATCCTCACCACGTAGGATCGTTGCTCGTC | 70 |
| oligo_24 [rn6]chrArc:845-875 | ACGTAGGATCGTTGCTCGTCACTCCCGTGGGTCCTCAAAGATCTGGGTATACGTAGGATCGTTGCTCGTC | 70 |
| oligo_25 [rn6]chrArc:891-921 | ACGTAGGATCGTTGCTCGTCTCAGAGCCACCCACCTGCCGCAGGTACTCTACGTAGGATCGTTGCTCGTC | 70 |
| oligo_26 [rn6]chrArc:922-952 | ACGTAGGATCGTTGCTCGTCGTGGTTCTGGATCTGGGACAGCCAATATTCACGTAGGATCGTTGCTCGTC | 70 |
| oligo_27 [rn6]chrArc:953-983 | ACGTAGGATCGTTGCTCGTCACTCCCACCACTTCTTGGCTGGCCCATTCAACGTAGGATCGTTGCTCGTC | 70 |
| oligo_28 [rn6]chrArc:984-1014 | ACGTAGGATCGTTGCTCGTCTCCACCCAGTTCTTCACCGAGCCCTGTTTGACGTAGGATCGTTGCTCGTC | 70 |
| oligo_29 [rn6]chrArc:1016-1046 | ACGTAGGATCGTTGCTCGTCCCTCACTGTACTGCAGAAACTCCTTCTTGAACGTAGGATCGTTGCTCGTC | 70 |
| oligo_30 [rn6]chrArc:1047-1077 | ACGTAGGATCGTTGCTCGTCTCCCGCTGAATGGCTTCGCGGGAGAGCGTAACGTAGGATCGTTGCTCGTC | 70 |
| oligo_31 [rn6]chrArc:1088-1118 | ACGTAGGATCGTTGCTCGTCACTGGTCAAGTGGCTCACCCTGCTTCTGTGACGTAGGATCGTTGCTCGTC | 70 |
| oligo_32 [rn6]chrArc:1119-1149 | ACGTAGGATCGTTGCTCGTCGTCTGGTACAGGTCCCGCTTACGCCAGAGGACGTAGGATCGTTGCTCGTC | 70 |
| oligo_33 [rn6]chrArc:1160-1190 | ACGTAGGATCGTTGCTCGTCCATACTGAATGATCTCCTCCTCCTCAGCGTACGTAGGATCGTTGCTCGTC | 70 |
| oligo_34 [rn6]chrArc:1204-1234 | ACGTAGGATCGTTGCTCGTCGTGGCGCAGAAAGCGCTTGAACTTGGGCTGACGTAGGATCGTTGCTCGTC | 70 |
| oligo_35 [rn6]chrArc:1249-1279 | ACGTAGGATCGTTGCTCGTCTTCCATGCCCCTCTGGATGAGCTGCTCCAGACGTAGGATCGTTGCTCGTC | 70 |
| oligo_36 [rn6]chrArc:1285-1315 | ACGTAGGATCGTTGCTCGTCAGAAGGCTCAGCTGCCTGCTCCAGGCCGTCACGTAGGATCGTTGCTCGTC | 70 |
| oligo_37 [rn6]chrArc:1316-1346 | ACGTAGGATCGTTGCTCGTCCAGTCTCATCCTCTGTGGGCAGAGGGGTGAACGTAGGATCGTTGCTCGTC | 70 |
| oligo_38 [rn6]chrArc:1347-1377 | ACGTAGGATCGTTGCTCGTCGACTCGCTGGTAAGAGCAGGCGTGAGTGCCACGTAGGATCGTTGCTCGTC | 70 |
| oligo_39 [rn6]chrArc:1378-1408 | ACGTAGGATCGTTGCTCGTCCTATTCAGGCTGGGTCCTGTCACTGGCTACACGTAGGATCGTTGCTCGTC | 70 |
| oligo_40 [rn6]chrArc:1434-1464 | ACGTAGGATCGTTGCTCGTCGACAAAAGCCACAGACTGGGTGTGGCAGGCACGTAGGATCGTTGCTCGTC | 70 |
| oligo_41 [rn6]chrArc:1465-1495 | ACGTAGGATCGTTGCTCGTCGTGTCAGCCCCAGCTCAATCAAGTCCTAGTACGTAGGATCGTTGCTCGTC | 70 |
| oligo_42 [rn6]chrArc:1496-1526 | ACGTAGGATCGTTGCTCGTCAGGTGTCTGGCTGGACAGGGCATCCCCTTGACGTAGGATCGTTGCTCGTC | 70 |
| oligo_43 [rn6]chrArc:1528-1558 | ACGTAGGATCGTTGCTCGTCGTGGCAGTTGTGAGTCAGGCCAGTGGGTGAACGTAGGATCGTTGCTCGTC | 70 |
| oligo_44 [rn6]chrArc:1560-1590 | ACGTAGGATCGTTGCTCGTCGGGGCTTCTTGATGTCCATGAATCATGGTTACGTAGGATCGTTGCTCGTC | 70 |
| oligo_45 [rn6]chrArc:1592-1622 | ACGTAGGATCGTTGCTCGTCGGGTAGGTGGCAGGTGGGAGCCCTATGGGAACGTAGGATCGTTGCTCGTC | 70 |
| oligo_46 [rn6]chrArc:1637-1667 | ACGTAGGATCGTTGCTCGTCGTGAGGCCACTGGAGACAGGGCCAGGACTAACGTAGGATCGTTGCTCGTC | 70 |
| oligo_47 [rn6]chrArc:1671-1701 | ACGTAGGATCGTTGCTCGTCAAAGGTGTTCTGTGATGGTCTGAGAGTGTAACGTAGGATCGTTGCTCGTC | 70 |
| oligo_48 [rn6]chrArc:1709-1739 | ACGTAGGATCGTTGCTCGTCCCAAAGGGCCCTGGACACTGATGCAGAATGACGTAGGATCGTTGCTCGTC | 70 |
| oligo_49 [rn6]chrArc:1755-1785 | ACGTAGGATCGTTGCTCGTCGGTGCCTACTTTTCATTGCCTTTCAGACACACGTAGGATCGTTGCTCGTC | 70 |
| oligo_50 [rn6]chrArc:1794-1824 | ACGTAGGATCGTTGCTCGTCTTCTGCTTTAGCATCTGCCCTGGGATGCCCACGTAGGATCGTTGCTCGTC | 70 |
| oligo_51 [rn6]chrArc:1844-1874 | ACGTAGGATCGTTGCTCGTCTAGGAGCGGGCTGCATCCCCGGAAGTAGAGACGTAGGATCGTTGCTCGTC | 70 |
| oligo_52 [rn6]chrArc:1875-1905 | ACGTAGGATCGTTGCTCGTCGGTAGGGCACCAGCTGGATCTGCTGTGTCTACGTAGGATCGTTGCTCGTC | 70 |
| oligo_53 [rn6]chrArc:1906-1936 | ACGTAGGATCGTTGCTCGTCGCCCAAGACTGGCCAGTTGCTCTGGGAGGCACGTAGGATCGTTGCTCGTC | 70 |
| oligo_54 [rn6]chrArc:1939-1969 | ACGTAGGATCGTTGCTCGTCTGCTTCAGCTCACCCTGAGAGGGGAGCTATACGTAGGATCGTTGCTCGTC | 70 |
| oligo_55 [rn6]chrArc:1970-2000 | ACGTAGGATCGTTGCTCGTCGGGCCAGGAGGCGCCAGCGCGTCAGGTCTGACGTAGGATCGTTGCTCGTC | 70 |
| oligo_56 [rn6]chrArc:2005-2035 | ACGTAGGATCGTTGCTCGTCGTCTGCTTTTCTTCACTGGTATGAATCACTACGTAGGATCGTTGCTCGTC | 70 |
| oligo_57 [rn6]chrArc:2036-2066 | ACGTAGGATCGTTGCTCGTCCCGCCTGGCATGGCTGAGTCATGGAGCCGAACGTAGGATCGTTGCTCGTC | 70 |
| oligo_58 [rn6]chrArc:2114-2144 | ACGTAGGATCGTTGCTCGTCCTGGTGTCATTCACCTGGCTCTGAAGACTCACGTAGGATCGTTGCTCGTC | 70 |
| oligo_59 [rn6]chrArc:2145-2175 | ACGTAGGATCGTTGCTCGTCTGGCCGGAAAGACTTCTCAGCAGCTTGAGAACGTAGGATCGTTGCTCGTC | 70 |
| oligo_60 [rn6]chrArc:2197-2227 | ACGTAGGATCGTTGCTCGTCTTCAAGAGAGGAGGGGACGGTGCTGGTGCTACGTAGGATCGTTGCTCGTC | 70 |
| oligo_61 [rn6]chrArc:2229-2259 | ACGTAGGATCGTTGCTCGTCTGAAAGTGTCTTGGAACCTCTGTGCAGGCAACGTAGGATCGTTGCTCGTC | 70 |
| oligo_62 [rn6]chrArc:2261-2291 | ACGTAGGATCGTTGCTCGTCACCTCCTCTTTGTAATCCTATTTTCTCTGCACGTAGGATCGTTGCTCGTC | 70 |
| oligo_63 [rn6]chrArc:2306-2336 | ACGTAGGATCGTTGCTCGTCACCTTCAGCTCTGAGGCTGAGCTGGGTGCTACGTAGGATCGTTGCTCGTC | 70 |
| oligo_64 [rn6]chrArc:2337-2367 | ACGTAGGATCGTTGCTCGTCCAGACCCGGGGTTTCACGCTGGCTTGTCTTACGTAGGATCGTTGCTCGTC | 70 |
| oligo_65 [rn6]chrArc:2368-2398 | ACGTAGGATCGTTGCTCGTCGGTGAGTGGCCAGCGGAGCGGGCATTCGTGACGTAGGATCGTTGCTCGTC | 70 |
| oligo_66 [rn6]chrArc:2409-2439 | ACGTAGGATCGTTGCTCGTCACAGACCCTGCTCAAGCTGCAGTGGCTTGTACGTAGGATCGTTGCTCGTC | 70 |
| oligo_67 [rn6]chrArc:2440-2470 | ACGTAGGATCGTTGCTCGTCCGCAGCGAACTGGGCTCTGTGCTGAGAGGGACGTAGGATCGTTGCTCGTC | 70 |
| oligo_68 [rn6]chrArc:2471-2501 | ACGTAGGATCGTTGCTCGTCCCTGCAAGGTTCTGGCGGGGGCCAAAGGCCACGTAGGATCGTTGCTCGTC | 70 |
| oligo_69 [rn6]chrArc:2502-2532 | ACGTAGGATCGTTGCTCGTCCAGGCTGGGCTAGGGCCCGAACCTTAAGGCACGTAGGATCGTTGCTCGTC | 70 |
| oligo_70 [rn6]chrArc:2537-2567 | ACGTAGGATCGTTGCTCGTCGACTTGACCAGCAGGCAGGGCACAGCAGGTACGTAGGATCGTTGCTCGTC | 70 |
| oligo_71 [rn6]chrArc:2570-2600 | ACGTAGGATCGTTGCTCGTCCTAGGAGCCAAGGCATGGGGTCTCCTGGGAACGTAGGATCGTTGCTCGTC | 70 |
| oligo_72 [rn6]chrArc:2606-2636 | ACGTAGGATCGTTGCTCGTCAATCACCCGGCAGGTCAGGGAAGTGCCTGGACGTAGGATCGTTGCTCGTC | 70 |
| oligo_73 [rn6]chrArc:2650-2680 | ACGTAGGATCGTTGCTCGTCACGAGGTGGTTGGTGCTGGGGTGTGGATGAACGTAGGATCGTTGCTCGTC | 70 |
| oligo_74 [rn6]chrArc:2683-2713 | ACGTAGGATCGTTGCTCGTCCTACTCAGACTACAGACACGAGCAGTTACCACGTAGGATCGTTGCTCGTC | 70 |
| oligo_75 [rn6]chrArc:2714-2744 | ACGTAGGATCGTTGCTCGTCGACCAGGCAGATGGAGGAACCTCAACATGGACGTAGGATCGTTGCTCGTC | 70 |
| oligo_76 [rn6]chrArc:2745-2775 | ACGTAGGATCGTTGCTCGTCCAGCAGTGGAACTGGTCTCAGAACACCAATACGTAGGATCGTTGCTCGTC | 70 |
| oligo_77 [rn6]chrArc:2781-2811 | ACGTAGGATCGTTGCTCGTCGGCTGGCAGGGGCACAGGGTGGGGGATCTGACGTAGGATCGTTGCTCGTC | 70 |

### Supplementary Table 1. Arc mRNA FISH probe

The table reports the RNA probes used in mRNA FISH experiment

| Imaging session | Marker | Concentration | Power at BFP | Number of frames |
|---|---|---|---|---|
| Dataset_1_day1 | mEGFP-Arc | 35 pM (R3) | 15 mW | 50000 (Pos0) |
| | | | | 30000 (Pos1) |
| | | | | 30000 (Pos2) |
| | | | | 23690 (Pos3) |
| | | | | 30000 (Pos4) |
| | CHC | 50 pM (R4) | 15 mW | 10000 |
| | PSD95 | 100 pM (R6) | 15 mW | 10000 |
| Dataset_1_day2 | mEGFP-Arc | 60 pM (R3) | 15 mW | 30000 |
| | CHC | 50 pM (R4) | 15 mW | 12500 |
| | PSD95 | 100 pM (R6) | 15 mW | 12500 |
| Dataset_2_CTRL_BrainPhys | mEGFP-Arc | 50 pM (R3) | 18 mW | 30000 |
| | CHC | 50 pM (R4) | 18 mW | 10000 |
| | PSD95 | 200 pM (R2) | 18 mW | 12000 |
| Dataset_2_CTRL_Charu | mEGFP-Arc | 50 pM (R3) | 18 mW | 30000 |
| | CHC | 30 pM (R4) | 18 mW | 15000 |
| | PSD95 | 150 pM (R2) | 15 mW | 15000 |

**Supplementary Table 2. DNA-PAINT imaging parameters**

The table reports the parameters used during the imaging acquisition of DNA-PAINT experiment.
